# TET2-dependent differential 5hmC deposition balances adult neural stem cell activation and differentiation

**DOI:** 10.64898/2025.12.12.693863

**Authors:** Laura Lázaro-Carot, Jordi Planells, Esteban Jiménez-Villalba, Leo J. Arteaga-Vázquez, Anjana Rao, Sacri R. Ferrón

## Abstract

Ten-eleven translocation (TET) enzymes are key regulators of active DNA demethylation, converting 5-methylcytosine (5mC) to 5-hydroxymethylcytosine (5hmC) and thereby shaping the epigenetic landscape and cellular identity. While their roles have been characterized in pluripotent and some multipotent stem cells, their function in adult neural stem cells (NSCs) of the subventricular zone (SVZ) remains poorly understood. Here, we show that TET2 is critical for the maintenance and differentiation of adult NSCs, orchestrating locus-specific 5hmC deposition across promoters, gene bodies, and enhancers. Importantly, 5hmC remodeling segregates into two functionally distinct programs: promoter-associated gains of 5hmC are strongly TET2-dependent and drive transcription of genes controlling neural differentiation and calcium signaling, whereas gene body- and enhancer-associated 5hmC gains partially depend on TET2 and sustain proliferative and metabolic pathways, thereby maintaining stemness. Loss of TET2 disrupts these 5hmC programs, downregulates key differentiation- and calcium-related genes, and impairs the normal differentiation-associated increase in intracellular calcium, revealing a functional consequence of altered epigenetic regulation. Together, our findings uncover a pivotal role for TET2 in coordinating complementary epigenetic and transcriptional programs that balance stemness and differentiation in adult NSCs.

## Introduction

Embryonic development is characterized by extensive neuronal production, whereas in adulthood, neurogenesis becomes restricted to discrete regions known as neurogenic niches. In vertebrates, two major neurogenic niches persist: the subgranular zone (SGZ) of the dentate gyrus (DG) in the hippocampus and the ventricular-subventricular zone (V-SVZ), lining the lateral ventricles (Kempermann et al., 2015; Lim & Alvarez-Buylla, 2016). Adult neurogenesis in these regions is sustained by the activity of neural stem cells (NSCs), which possess the capacity for self-renewal and multipotent differentiation into neurons, astrocytes and oligodendrocytes (Belenguer et al., 2016; Suh et al., 2007). NSCs are evolutionarily related to developmental radial glia and share astroglial features, including expression of glial fibrillary acidic protein (GFAP) and glutamate-aspartate transporter (GLAST) (Cebrian-Silla et al., 2021; Doetsch, 2003; Llorens-Bobadilla et al., 2015). Among both niches, the V-SVZ represents the most active site of neurogenesis in the adult brain (Obernier & Alvarez-Buylla, 2019). Upon activation, V-SVZ NSCs generate transit-amplifying progenitor (TAPs) that give rise to neuroblasts (Obernier et al., 2018), which migrate through the rostral migratory stream (RMS) to the olfactory bulb (OB), where they differentiate into functional neurons involved in olfactory processing (Obernier & Alvarez-Buylla, 2019). In addition, V-SVZ-derived NSCs also produce oligodendrocytes and astrocytes that migrate to the corpus callosum and striatum, respectively (Delgado et al., 2021; Figueres-Oñate et al., 2019; Sohn et al., 2015).

Communication between NSCs and their surrounding microenvironment is essential for the regulation of adult neurogenesis (Chaker et al., 2016; Urbán et al., 2019). This dialogue integrates extrinsic niche-derived signals with intrinsic transcriptional programs within NSCs through DNA regulatory elements such as enhancers and promoters that serve as hubs for the convergence of transcription factors and epigenetic modifiers (Dall’Agnese & Young, 2023). Epigenetic regulation thus plays a fundamental role in the spatial and temporal control of lineage-specific gene expression (Friedman et al., 2024; Wagh et al., 2025). The primary epigenetic modifications including DNA methylation, histone modifications, and non-coding RNAs, that collectively remodel the chromatin landscape determining cell identity (Atlasi & Stunnenberg, 2017; Honig & Murrell, 2025).

Although 5-methylcytosine (5mC) was regarded as a stable and irreversible covalent DNA modification, the discovery of ten-eleven translocation (TET) enzymes, which catalyze the oxidation of 5mC to 5-hydroxymethylcytosine (5hmC) (Ko et al., 2010; Tahiliani et al., 2009) has fundamentally revised this view, revealing the dynamic nature of cytosine methylation. TET enzymes can further oxidize 5hmC to 5-formylcytosine (5fC) and 5-carboxylcytosine (5caC), both of which can be removed through base excision repair involving the DNA repair enzyme thymine DNA glycosylase (He et al., 2011; Ito et al., 2010). Genome-wide studies have shown that 5hmC accumulates at open chromatin regions, gene bodies of highly expressed genes, and cis-regulatory elements such as enhancers and promoters, where TET enzymes modulate transcriptional activity (Bogdanović et al., 2016; Huang et al., 2014; Tsagaratou et al., 2014). However, the regulatory role of 5hmC is context-dependent, acting as either a permissive or a repressive mark depending on the chromatin environment (Palczewski et al., 2025). Thus, 5hmC constitutes a dynamic epigenetic regulatory layer that integrates DNA methylation with transcriptional control (Li et al., 2019; Rasmussen et al., 2019; Xu et al., 2011).

The brain is highly enriched in 5hmC compared to other adult tissues (Globisch et al., 2010; Kriaucionis & Heintz, 2009; Münzel et al., 2010; Szwagierczak et al., 2010). Moreover, 5hmC levels increase during postnatal maturation and aging (Sun et al., 2014; Szulwach et al., 2011; Wen et al., 2014), and its distribution varies across brain regions and developmental stages, emphasizing its importance in neural function (Wu & Zhang, 2017). Consistently, TET proteins have been implicated in neuronal differentiation and synaptic plasticity through the regulation of neurogenic transcriptional programs (Guo et al., 2011; X. Li et al., 2017; Lian et al., 2016; Zhang et al., 2013). However, the contribution of TET proteins to the regulation of adult NSCs within the V-SVZ neurogenic niche remains largely unknown.

In this study, we uncover a key role of TET2 in regulating adult NSC dynamics in the V-SVZ. Using complementary *in vivo* and *in vitro* approaches, we show that loss of *Tet2* disrupts the NSC pool, resulting in impaired adult neurogenesis and gliogenesis. *Tet2*-deficient NSCs display enhanced self-renewal and proliferative capacity, accompanied by a pronounced reduction in differentiation potential *in vitro*. We further demonstrate that TET2 modulates 5hmC deposition at enhancers, promoters, and gene bodies during NSC differentiation, and that these region-specific 5hmC changes correlate with transcriptional alterations. Notably, the transcriptional programs associated with 5hmC are distinct across regulatory regions: enhancer-associated 5hmC is primarily linked to the maintenance of the stem cell state, promoter-associated 5hmC is enriched at genes controlling calcium signaling and homeostasis, and gene-body 5hmC correlates with pathways involved in differentiation, calcium function, and aspects of stemness. Together, these findings indicate that TET2 orchestrates distinct regulatory mechanisms across genomic contexts, and that the integration of these mechanisms fine-tunes the balance between self-renewal and differentiation in adult NSCs within the V-SVZ niche, revealing a previously unrecognized role for TET2 in coupling epigenetic control with functional outcomes in neural stem cell biology.

## Results

### TET2 is expressed in adult NSCs from the V-SVZ and increases upon differentiation

TET expression in adult neurogenic niches has been previously reported (X. Li et al., 2017; Montalbán-Loro et al., 2019). Transcriptional analysis confirmed that *Tet2* is highly expressed in adult brain regions, including the V-SVZ (**Figure 1A**). In the presence of fibroblast growth factor (FGF) and epidermal growth factor (EGF), V-SVZ-derived NSCs proliferate and form multipotent clonal aggregates known as neurospheres (Jiménez-Villalba et al., 2024). To induce neural progenitor differentiation, neurospheres are transferred to adherent conditions and cultured for 2 days in FGF-containing medium (2 DIV; **Figure 1A**) followed by mitogen withdrawal and serum addition to trigger terminal differentiation into neurons, oligodendrocytes, and astrocytes (7 DIV in total) (**Figure 1A**) (Belenguer et al., 2016). Although expressed at lower levels in proliferating V-SVZ-derived neurospheres, *Tet2* is markedly upregulated during *in vitro* differentiation, showing the most robust induction among TET family members (**Figure 1A, B**). Notably, *Tet2* is also highly expressed in terminally differentiated astrocytes (**Figure 1A**). The murine *Tet2* gene generates two alternatively spliced isoforms (**Figure S1A**), but their expression patterns in brain cells remain poorly defined. Isoform-specific qPCR revealed that isoform 1 is the predominant variant in both whole brain and in adult V-SVZ-derived NSCs, in contrast to its expression profile in non-neural tissues (**Figure S1A**).

**Figure 1.**
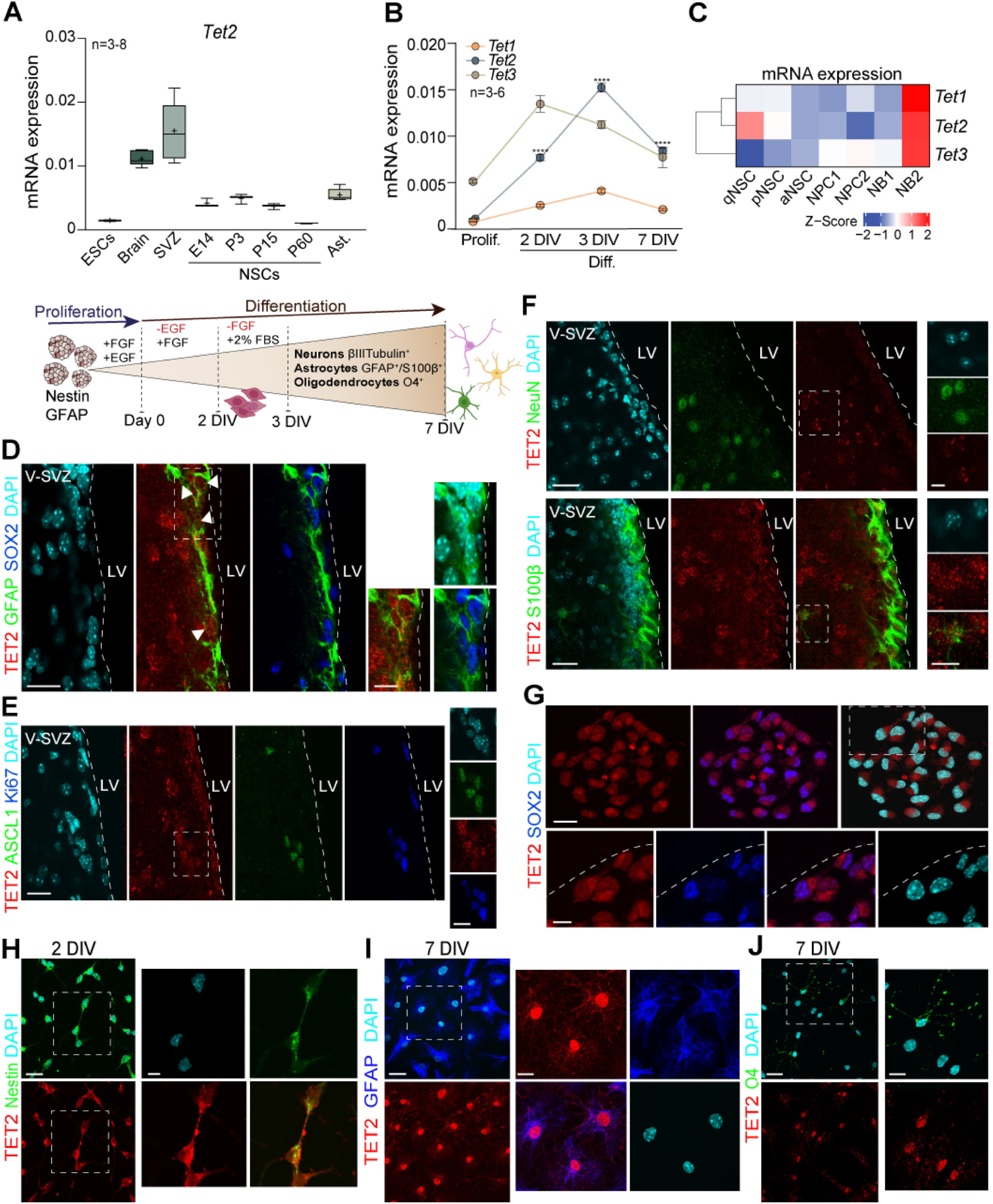
*Tet2* is expressed in adult NSCs and increases during the differentiation process. **A)** qPCR analysis of *Tet2* expression across various cell types and tissues (upper panel). Schematic of the *in vitro* differentiation protocol (lower panel). Adult NSCs were expanded under proliferating conditions with EGF and FGF. Differentiation was initiated by EGF withdrawal and plating on matrigel. After two days *in vitro* (2 DIV), FGF was replaced with 2% fetal bovine serum (FBS) and cells were fixed for immunostaining after 7 days (7 DIV). **B)** qPCR analysis of *Tet1*, *Tet2,* and *Tet3* expression in proliferating and differentiating adult NSCs after 2, 3, and 7 DIV. **C)** Heatmap of RNA-seq expression levels for murine *Tet* family genes in V-SVZ-sorted quiescent NSCs (qNSC), primed NSCs (pNSCs) and activated NSCs (aNSCs). Neural progenitors (NPC) and neuroblast (NB) populations are also shown. Data obtained from gene expression omnibus (GEO) with accession GSE138243. **D-F)** Immunohistochemistry images for TET2 (red, **D-F**), GFAP (green) and SOX2 (blue) (**D**), ASCL1 (green) and Ki67 (blue) (**E**), and NeuN (green) and S100β (green) (**F**) in the adult V-SVZ of wild-type mice. **H)** Immunocytochemistry images for TET2 (red) and Nestin (green) in NSCs after 2 days of differentiation. **I-J)** Immunocytochemistry images for TET2 (red, **I-J**), and GFAP (blue, **I**) and O4 (green, **J**) in NSCs after 7 days of differentiation. *Gapdh* was used to normalize qPCR data. DAPI was used to counterstain DNA. Ast.: astrocytes; E: embryonic day P: postnatal day; V-SVZ: ventricular-subventricular zone; LV: lateral ventricle. In box-and-whiskers plots, the mean is indicated by (+) and whiskers denote minimum and maximum values. In B, mean ± s.e.m. are shown. Statistical significance was evaluated using ANOVA tests. P-values and sample size (n) are indicated. ****: P<0.0001. Scale bars in D-G: 10 μm (high-magnification images, 5 μm); in H-J: 10 μm (high-magnification images in H, I: 7 μm; in J: 5 μm). See also Figure S1.

In the adult V-SVZ, NSCs exist in multiple activation states, which are tightly regulated. Three NSC subpopulations can be distinguished based on GLAST and EGF receptor (EGFR) expression: quiescent NSCs (qNSCs, GLAST⁺/EGFR⁻), primed-for-activation NSCs (pNSCs, GLAST⁺/EGFR^low^), and activated NSCs (aNSCs, GLAST⁺/EGFR⁺) (Belenguer et al., 2021). Analysis of previously published RNA-seq datasets from our group revealed that *Tet2* expression is particularly enriched in qNSCs and late neuroblasts (**Figure 1C**, NB2), suggesting that TET2 may act at both early and late stages of the neurogenic lineage, potentially contributing to the maintenance of NSC quiescence as well as to later steps of neuronal differentiation.

At the protein level, TET2 was detected in a subset of GFAP^+^/SOX2^+^ NSCs lining the lateral ventricles (**Figures 1D** and **S1B, C**). As cells progressed along the neurogenic lineage toward ASCL1^+^ transit-amplifying progenitors (TAPs), TET2 appeared increasingly enriched within the nucleus (**Figure 1E**). This upregulation was further evident in differentiated progeny within the striatal parenchyma, including NeuN^+^ neurons and S100β^+^ astrocytes (**Figure 1F**). Likewise, cells within the olfactory bulb (OB), the primary destination of V-SVZ-derived neuroblasts, exhibited robust nuclear TET2 expression (**Figure S1D**). Consistent with these *in vivo* observations, TET2 was also present in SOX2^+^ neurosphere cultures (**Figure 1G),** with higher expression levels observed in Nestin^+^ progenitors and in GFAP^+^ astrocytes during *in vitro* differentiation (**Figure 1H-J**). Together, these findings support the presence of TET2 across the adult V-SVZ neurogenic lineage and suggest a potential role for TET2 in regulating NSC proliferation and lineage commitment during adult neurogenesis.

### TET2 balances neurogenesis and gliogenesis in the adult V-SVZ

To investigate the role of TET2 in the regulation of adult V-SVZ NSCs, we generated a conditional *Tet2* knockout mouse model by crossing *Tet2^loxP/loxP^* males (Moran-Crusio et al., 2011) with female mice expressing Cre recombinase under the control of the *Gfap* promoter (*Gfap-Cre^+/o^*) (Garcia et al., 2004) (**Figure S2A**). This strategy results in the deletion of exon 3 in GFAP-expressing cells, including adult NSCs and astrocytes, leading to production of a truncated, non-functional TET2 protein (**Figure S2A**). *Cre* expression was confirmed in the brain and in V-SVZ-derived NSCs from *Tet2^l^*^oxp/loxp^-*Gfap-Cre*^+/0^ (*Tet2-Gfap^cre^*) mice, but not in *Tet2^loxP/loxp-^Cre^0/0^* controls (*Tet2-Gfap^control^*) (**Figure S2B)**. Consistent with efficient recombination, *Tet2* mRNA and protein expression were selectively reduced in NSCs and astrocytes of *Tet2-Gfap^cre^* mice, without compensatory changes in *Tet1* and *Tet3* transcript levels (**Figures 2A** and **S2C, D**). Body and brain weights were comparable between genotypes (**Figure S2E**), indicating that *Tet2* deletion does not alter general growth or gross brain morphology.

**Figure 2.**
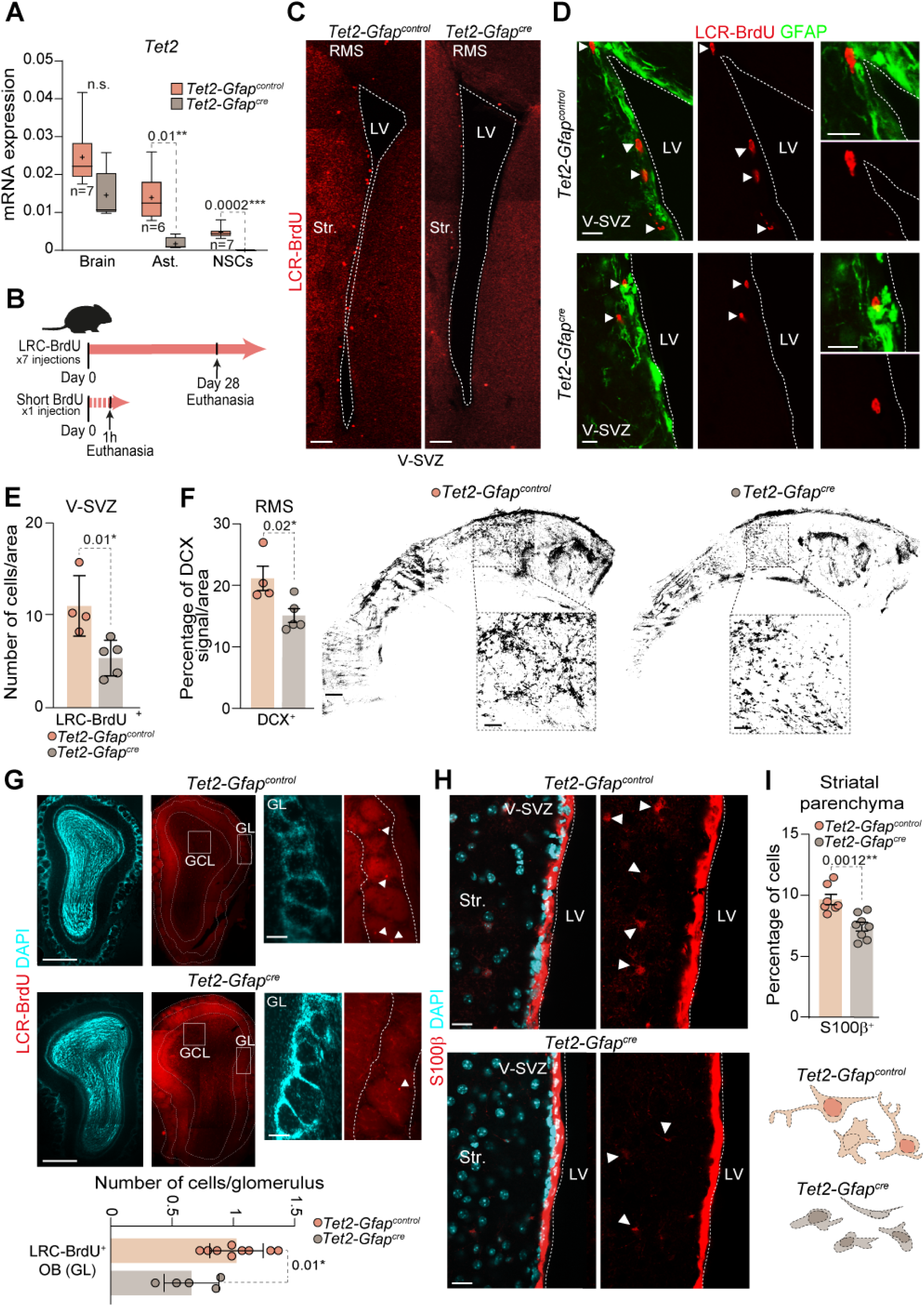
*Tet2* deletion disrupts NSCs homeostasis *in vivo*. **A)** qPCR analysis of *Tet2* expression in whole brain, astrocytes (Ast.) and NSCs isolated from the adult V-SVZ from *Tet2-Gfap^control^* and *Tet2-Gfap^cre^* mice. **B)** Schematic of the BrdU paradigms illustrating both long-term label-retaining cell (LRC) and short-term labeling protocols. **C)** Panoramic immunohistochemistry of the V-SVZ from *Tet2-Gfap^control^* and *Tet2-Gfap^cre^* mice showing BrdU^+^ LRCs (red). **D)** High-magnification images of BrdU^+^ LRCs (red) and GFAP (green) in the V-SVZ of both genotypes. **E)** Quantification of BrdU^+^ LRCs in the V-SVZ of *Tet2-Gfap^control^* and *Tet2-Gfap^cre^* mice. **F)** Percentage of doublecortin (DCX) signal per area in the RMS of both genotypes (left panel). Confocal whole-mount images of DCX staining in the RMS (right panel). **G)** BrdU^+^ LRCs (red) in the glomerular layer (GL) of the olfactory bulb (OB) of *Tet2-Gfap^control^* and *Tet2-Gfap^cre^* mice (upper panel). Quantification of newly generated neurons in the GL (lower panel). **H)** Immunohistochemistry images for S100β^+^ differentiated astrocytes (red) in the striatal parenchyma (Str.) of both genotypes. **I)** Quantification of S100β^+^ cells as a percentage of total DAPI^+^ nuclei in the striatal parenchyma of *Tet2-Gfap^control^* and *Tet2-Gfap^cre^* mice (upper panel). Representative morphological reconstructions of S100β^+^ astrocytes (lower panel). DAPI was used to counterstain DNA. *Gapdh* served as the normalization control for qPCR analyses. V-SVZ: ventricular-subventricular zone; LV: lateral ventricle; RMS: rostral migratory stream. In box-and-whiskers plots, the mean is indicated by (+) and whiskers denote minimum and maximum values. In bar plots, data represent mean ± s.e.m. Statistical significance was assessed using unpaired two-tailed t tests and Mann–Whitney tests. P values and sample sizes (n; individual data points shown) are indicated. *: P<0.05; **: P<0.01; ***: P<0.001; n.s.: not significant. Scale bars in C 100 μm; in D: 5 μm; in G: 500 μm (high-magnification images, 100 μm); in H: 10 μm. See also Figure S2.

To assess NSC dynamics *in vivo*, two-month-old *Tet2-Gfap^cre^* and *Tet2-Gfap^control^* mice were injected with BrdU 28 days prior to analysis (**Figure 2B**). BrdU is retained in slowly dividing, label-retaining cells (LRCs), which include quiescent NSCs and newborn neurons that exit cell cycle upon differentiation. The number of slowly dividing LRCs in the V-SVZ was significantly reduced in *Tet2*-deficient mice (**Figure 2C-E**), indicating a potential decrease of the NSC pool following *Tet2* deletion. Flow cytometry analysis confirmed a reduced proportion of total NSCs in *Tet2*-deficient mice, consistent with the BrdU retention results (**Figure S2F**). Notably, besides the overall decline in the number of NSCs, we observed a shift in NSC subpopulation composition, characterized by a marked decrease in GLAST⁺/EGFR⁻quiescent NSCs and a corresponding increase in GLAST⁺/EGFR^low^ primed-for-activation NSCs (**Figure S2F**). These data support a role for TET2 in sustaining both the size and compositional balance of the adult NSC pool.

Despite the reduction in NSCs, overall proliferative activity in the niche remained unchanged in *Tet2-Gfap^cre^* mice, as evidenced by comparable Ki67 immunoreactivity and short-term BrdU incorporation (1h before sacrifice) across genotypes (**Figure 2B** and **S2G, H**). Similarly, the proportion of proliferating ASCL1^+^ progenitors and slowly dividing LRCs that were Ki67^+^ was unaffected (**Figure S2G, I, J**). Functionally, the reduction in NSCs was associated with a significant decrease in doublecortin-positive (DCX^+^) neuroblasts and a sparser rostral migratory stream in *Tet2-Gfap^cre^* mice (**Figure 2F**). Consistently, the number of BrdU^+^ newborn neurons in the glomerular layer (GL) of the OB was significantly reduced (**Figure 2G**), likely reflecting impaired neurogenesis from the *Tet2*-deficient V-SVZ. This neurogenic deficit coincided with a reduced proportion of terminally differentiated S100β^+^ astrocytes in the striatal parenchyma of *Tet2-Gfap^cre^* mice (**Figure 2H, I**). In contrast, no significant differences were detected in the density of newly formed oligodendrocytes in the corpus callosum (CC) (**Figure S2K**) or in the number of BrdU^+^ neurons in the granule cell layer of the OB (**Figure S3L**). Together, these findings indicate that TET2 is required to maintain the adult NSC pool and to preserve the neurogenic-gliogenic balance in the V-SVZ *in vivo*.

### TET2 promotes terminal astrocytic differentiation of NSCs from the adult V-SVZ

To further elucidate how TET2 regulates adult NSC function, V-SVZ tissue from 2-4 month-old *Tet2-Gfap^control^* and *Tet2-Gfap^cre^* mice was isolated, dissociated, and cultured under proliferation-promoting neurosphere conditions. *Tet2-*deficient V-SVZs generated a greater number of primary neurospheres compared to controls (**Figure 3A, B**). To assess self-renewal, primary neurospheres were dissociated to single cells and plated at limiting dilution across sequential passages. *Tet2*-deficient cultures consistently produced more secondary neurospheres with an elevated proliferative rate as a higher proportion of Ki67^+^ cells within *Tet2-Gfap^cre^* neurospheres was observed (**Figure 3A-C**). Accordingly, *Tet2-Gfap^cre^* neurospheres contained a higher number of cells per sphere (**Figure S3A**) and formed larger secondary neurospheres (**Figure S3B**), resulting in a greater cumulative expansion over successive passages in *Tet2-*deficients cultures (**Figure 3D**). Importantly, apoptosis rates were comparable between genotypes (1.44 ± 0.31 in *Tet2-Gfap^control^ vs*. 1.30 ± 0.14 in *Tet2-Gfap^cre^*), indicating that increased neurosphere formation was not attributable to reduced cell death.

**Figure 3.**
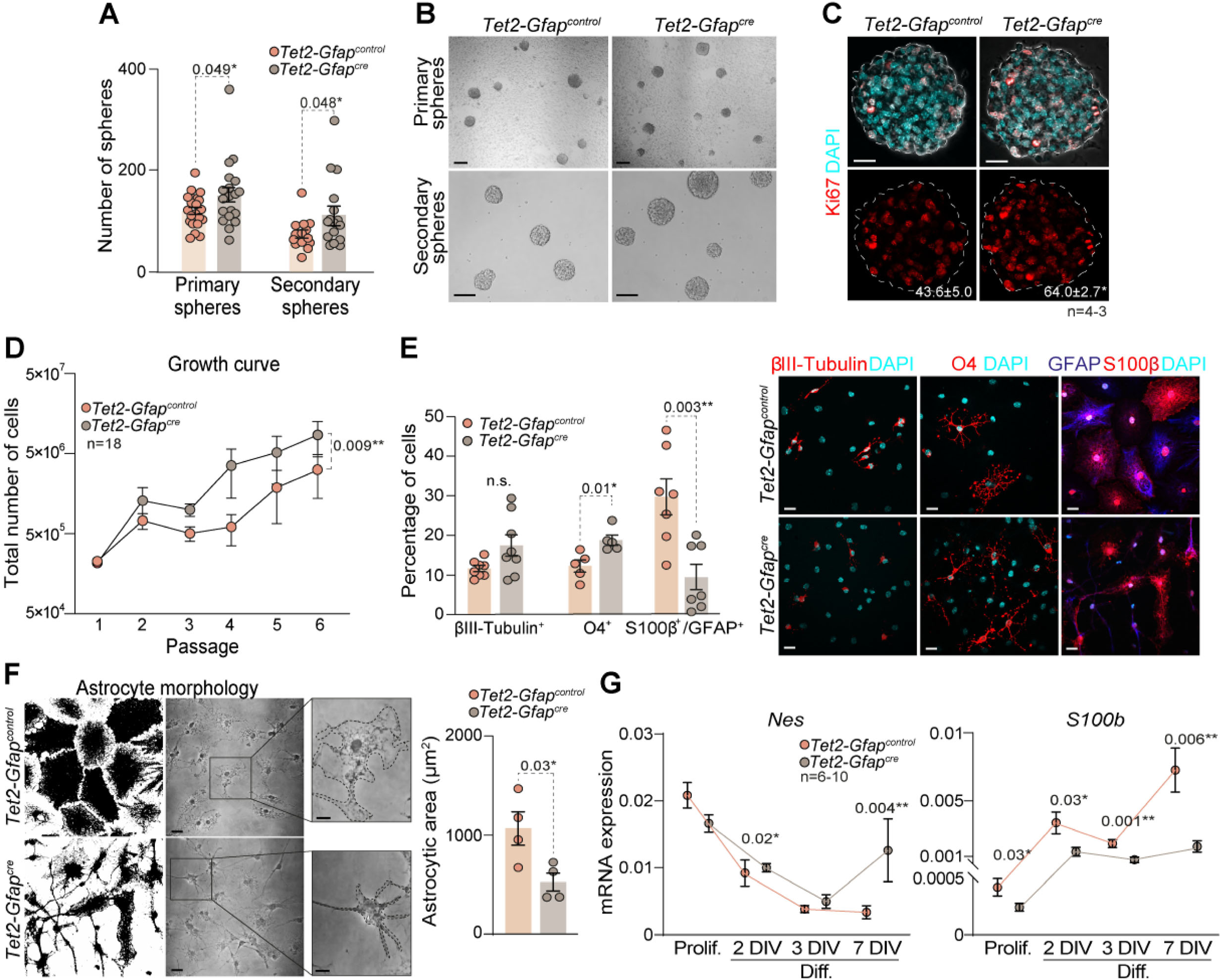
Loss of *Tet2* impairs astrocytic terminal differentiation of adult V-SVZ NSCs. **A)** Quantification of primary and secondary neurosphere formation from V-SVZ-derived NSCs of *Tet2-Gfap^control^* and *Tet2-Gfap^cre^* mice. **B)** Representative phase-contrast images of primary and secondary neurospheres generated from both genotypes. **C)** Immunocytochemistry images of Ki67 (red) in secondary neurospheres from adult wild-type V-SVZ-derived NSCs. Quantification of Ki67^+^ proliferating cells as a percentage of total DAPI^+^ cells per neurosphere is indicated. **D)** Growth curve showing total cell numbers across six passages in neurosphere cultures derived from *Tet2-Gfap^control^* and *Tet2-Gfap^cre^* mice. **E)** Percentage of βIII-Tubulin*^+^* neurons, O4^+^ oligodendrocytes and GFAP^+^/S100β*^+^* astrocytes among total DAPI^+^ cells in both genotypes after 7 DIV of differentiation (left panel). Representative immunocytochemistry images of differentiated NSCs stained for βIII-Tubulin (green), O4 (red), GFAP (blue), and S100β (red) after 7 DIV of differentiation (right panel). **F)** Binary confocal images (left) and phase-contrast images (right) of differentiated astrocytes from both genotypes at 7 DIV (left panel).Quantification of astrocytic cell area (μm^2^) at 7 DIV in cultures from *Tet2*-*Gfap^control^* and *Tet2*-*Gfap^cre^* mice (right panel). **G)** qPCR analysis of the undifferentiation marker *Nes* and the astrocytic differentiation marker *S100b* in NSCs under proliferation (Prolif.) and during in vitro differentiation (2, 3 and 7 DIV). DAPI was used to counterstain DNA. *Gapdh* was used to normalize qPCR data. DIV: days *in vitro*. The mean and the s.e.m are indicated in plots. Significance was evaluated using linear regression, unpaired two-tailed t-test and Mann-Whitney test. P-values and number of samples (n and dots) are indicated. *: P<0.05; **: P<0.01; n.s.: not significant. Scale bars in B: 200 μm (top), 100 μm (bottom); in C: 15 μm; in E, F: 10 μm (high-magnification: 5 μm). See also Figure S3.

Adult NSCs from the V-SVZ are multipotent and generate neurons, astrocytes and oligodendrocytes *in vitro*, recapitulating their *in vivo* lineage potential (Jiménez-Villalba et al., 2024). To evaluate lineage specification, *Tet2-Gfap^control^* and *Tet2-Gfap^cre^* neurospheres were induced to differentiate (**Figure 1A**). After seven days in differentiation conditions, *Tet2*-deficient cultures displayed a significant reduction in GFAP^+^/S100β^+^ mature astrocytes (**Figure 3E**), and these astrocytes exhibited altered filament organization and reduced surface area (**Figure 3F**). Conversely, the proportion of O4^+^ oligodendrocytes was increased in *Tet2-Gfap^cre^* cultures (**Figure 3E**), consistent with lineage biases reported in TET-deficient haematopoietic cells (An et al., 2015; Moran-Crusio et al., 2011; Tsagaratou et al., 2017).

qPCR analysis showed that expression of the progenitor marker *Nestin (Nes),* which normally declines during differentiation, remained elevated in *Tet2-Gfap^cre^* cultures (**Figure 3G**). In contrast, the mature astrocytic marker *S100b*, which normally increases during differentiation, was significantly reduced in *Tet2*-deficient cultures at both 2 and 7 days *in vitro* (**Figure 3G**). These molecular profiles were consistent with a less differentiated cellular state. This interpretation was further supported by increased Ki67^+^ cell frequency during differentiation in *Tet2-*deficient cultures (**Figure S3C**). Expression of neuronal (*Tubb3*) and oligodendroglial (*Olig2)* lineage markers remained unchanged between genotypes (**Figure S3D**). Together, these findings indicate that loss of *Tet2* maintains NSCs in a more undifferentiated state and impairs astrocytic maturation, supporting a crucial role for TET2 in promoting astroglial lineage commitment in the adult V-SVZ.

### Deletion of *Tet2* alters gene expression programs during adult NSC differentiation

To investigate the molecular mechanisms underlying the impaired proliferation and differentiation of *Tet2*-deficient NSCs, we performed RNA-seq on proliferating and early differentiated NSCs derived from *Tet2-Gfap^control^* and *Tet2-Gfap^cre^* mice. Hierarchical clustering of differentially expressed genes (DEGs) revealed robust transcriptional remodeling during differentiation in both genotypes, consistent with global transition in cellular state (**Figure 4A** and **S4A**). Comparative analysis identified 4,932 genes upregulated and 5,535 genes downregulated during differentiation in both wild-type and *Tet2*-deficient cells (**Figure 4A**). However, 4,136 genes were differentially regulated between genotypes, indicating a *Tet2*-dependent transcriptional module required for proper NSC lineage progression (**Figure 4A** and **S4A**). To clarify stage-specific effects of *Tet2* deletion, we analyzed gene expression changes separately in proliferating and differentiated cultures. Under proliferative conditions, loss of *Tet2* altered the expression of 623 genes, whereas under differentiation conditions, 2,131 genes were dysregulated (**Figure 4B** and **S4B**), representing more than a threefold increase in transcriptional dysregulation (**Figure 4B** and **S4B**). These findings indicate that TET2 plays a relatively important regulatory role in proliferating NSCs that becomes critical during differentiation.

**Figure 4.**
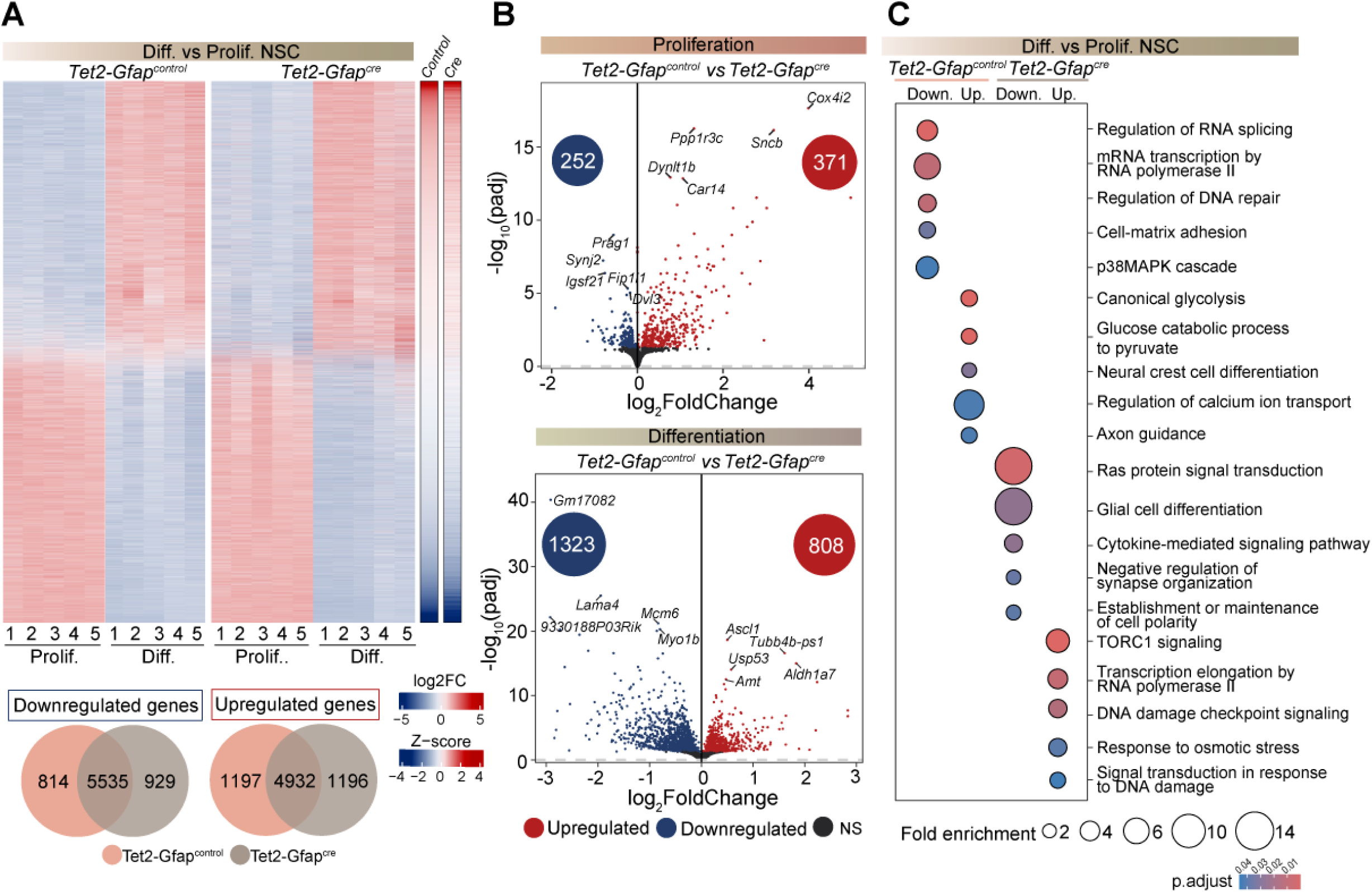
TET2 orchestrates the transcriptomic regulation of adult NSCs. **A)** Heatmap showing Z-score-scaled expression of all differentially expressed genes (DEGs) identified in *Tet2-Gfap^control^* and *Tet2-Gfap^cre^* NSCs cultures across the differentiation process (top panel). Log (fold change) in gene expression levels in proliferation (Prolif.) vs 3 days of differentiation (Diff.) conditions of NSCs from both genotypes is also indicated. Venn diagrams summarizing the number of DEGs shared or uniquely regulated between genotypes (bottom panel). **B)** Volcano plots illustrating RNA-seq differential expression analysis results (log_2_ fold change) comparing *Tet2-Gfap^control^* and *Tet2-Gfap^cre^* NSCs under proliferative conditions (top panel) and after 3 days of *in vitro* differentiation (botom panel). Upregulated genes are shown in red and downregulated genes in blue. Non-significant (NS) changes are shown in black. **C)** Gene ontology (GO) enrichment analysis of biological processes associated with genes upregulated or downregulated in *Tet2-Gfap^control^* and *Tet2-Gfap^cre^* NSCs during differentiation. See also Figure S4.

Gene ontology (GO) enrichment analysis showed that wild-type NSCs activated developmental and maturation-associated programs during differentiation, including axon guidance and calcium signalling regulation (**Figure 4C** and **S4B**). In contrast, *Tet2*-deficient NSCs failed to induce these lineage-specific pathways and instead exhibited aberrant enrichment of stress-associated programs, such as DNA damage response and osmotic stress signaling (**Figure 4C** and **S4B**), suggesting a shift from differentiation toward a stress-adaptative state. Additionally, *Tet2*-deficient cultures displayed reduced expression of essential processes for neural maturation, including glial cell differentiation, synapse organization, and maintenance of cell polarity (**Figure 4C** and **S4B**), consistent with their impaired differentiation phenotype. Notably, gene programs that are normally downregulated during wild-type NSC differentiation, such as RNA splicing, transcriptional regulation, DNA repair, and p38 MAPK signalling, remained inappropriately sustained in *Tet2*-deficient cells (**Figure 4C**). Together, these results indicate that TET2 is required not only for the activation of lineage-specific differentiation programs but also for the timely repression of core proliferative and stress-associated gene networks that must be silenced to enable adult NSC differentiation.

### Differential TET2-dependent 5hmC deposition at regulatory elements balances stemness maintenance and differentiation in adult NSCs

Given that TET2 modulates transcriptional programs in adult NSCs, we first examined whether this function depends on its DNA hydroxymethylation activity. Immunofluorescence analysis revealed a marked reduction in global 5hmC levels in *Tet2*-deficient NSCs compared with controls (**Figure S5A**). To map genome-wide 5hmC distribution, we performed cytosine-5-methylenesulfonate immunoprecipitation (CMS-IP) followed by high-throughput sequencing in proliferating and 3 DIV-differentiating wild-type and *Tet2*-deficient NSCs (Huang et al., 2012). Genome-wide 5hmC profiling showed broad distribution in proliferating NSCs, with enrichment at promoters, exons and introns (**Figure S5B**). Upon differentiation, global 5hmC levels increased substantially, coinciding with elevated TET2 expression (**Figure S5B**), and gains were predominantly observed at promoters, gene bodies and enhancers (**Figure S5B**), as previously observed in other tissues (Bai et al., 2025; He et al., 2021; Tsagaratou et al., 2014). Integration of CMS-IP and RNA-seq data revealed a positive association between 5hmC accumulation and transcriptional activity, with highly expressed genes exhibiting the strongest enrichment (**Figure S5C**). Consistent with observations in other cell types (Cui et al., 2020; Tsagaratou et al., 2014), 5hmC was depleted near transcription start sites (TSSs) of highly and moderately expressed genes, whereas lowly expressed genes showed the opposite pattern (**Figure S5C, D**). Importantly, differentiation-linked increases in 5hmC at promoters, gene bodies, and enhancers were largely TET2-dependent, as *Tet2* loss markedly reduced deposition at these regions (**Figure 5A** and **S5B-D**).

**Figure 5.**
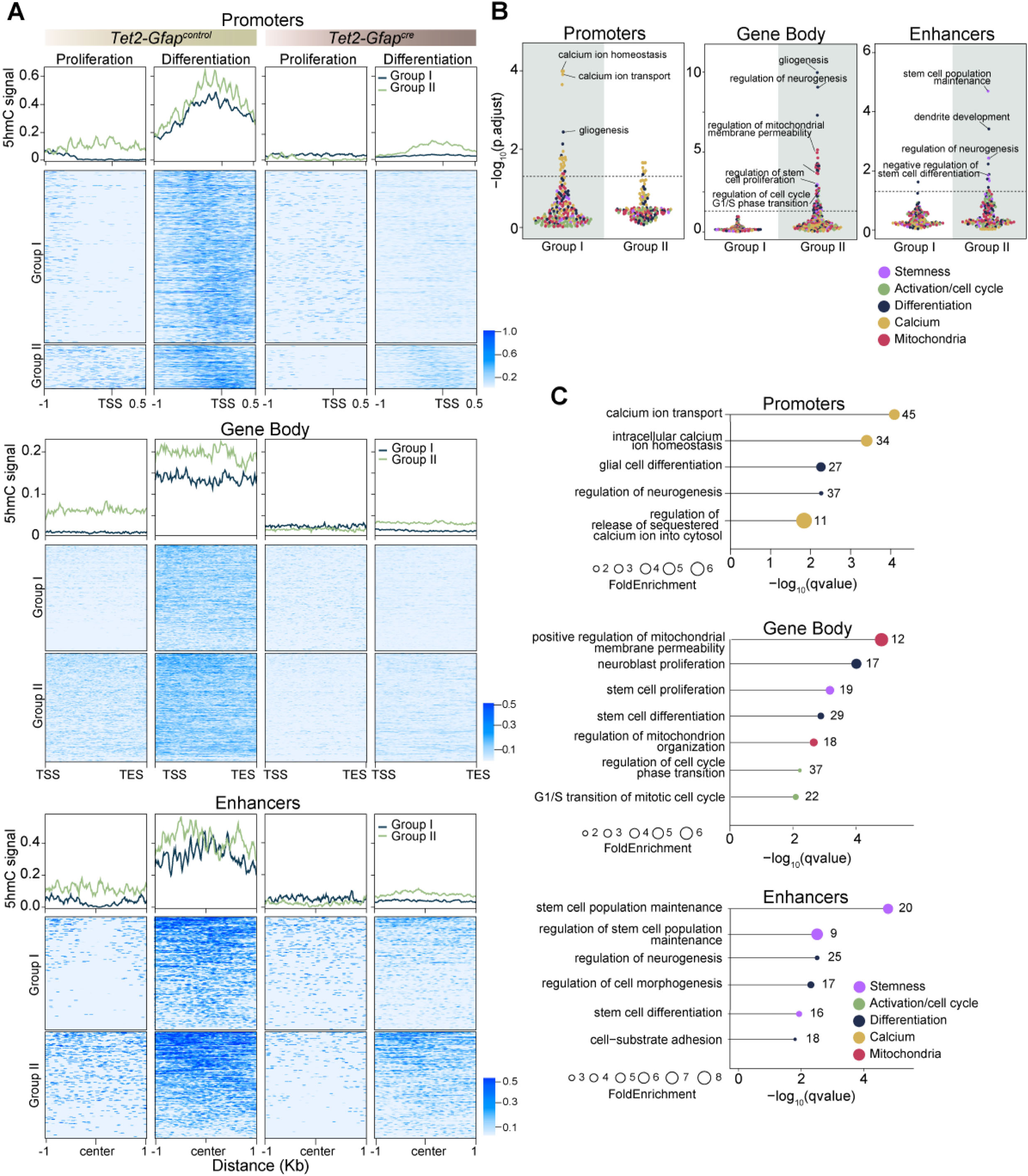
5hmC deposition mediated by TET2 reveals distinct regulatory roles at promoters, gene body, and enhancers during adult NSCs differentiation. **A)** Metagenes (top) and heatmaps (bottom) displaying genes that gain 5hmC during NSCs differentiation at promoters (upper panel), gene bodies (middle panel), and enhancers (lower panel) in *Tet2*-*Gfap^control^ and Tet2*-*Gfap^cre^* cultures. Genes that gain 5hmC during differentiation were separated into Group I and Group II based on TET2 dependency. **B)** Jitter plots of associated gene ontology (GO) terms enriched in genes from Group I and II of promoters, gene bodies and enhancers. Dashed lines indicate p-adj<0.05. **C)** Bar plots of selected significantly enriched biological processes associated with genes from Group I and Group II of the different analyzed genomic features. Each GO category is indicated by a color. Dot size represents fold enrichment. Number of genes per category is indicated. X-axis represents –log_10_(q-value). See also Figure S5.

To identify genomic regions requiring TET2 for hydroxymethylation, we compared *loci* that gained 5hmC in wild-type versus *Tet2-*deficient NSCs. This analysis delineated two groups of features. Group I gained strong 5hmC at promoters, gene bodies, or enhancers during differentiation in wild-type cells but showed minimal or no gains in the absence of TET2, indicating strict dependence of TET2 (**Figure 5A**). In contrast, Group II exhibited elevated basal 5hmC levels in proliferating NSCs, primarily within gene bodies and enhancers, and further increased 5hmC upon differentiation (**Figure 5A**). However, *Tet2*-deficient NSCs failed to reach the wild-type levels (**Figure 5A**), demonstrating that TET2 is partially required for achieving the full hydroxymethylation state associated with differentiation.

Next, we examined whether these region-specific 5hmC patterns correlated with distinct biological processes. GO-terms enrichment analyses revealed that effects due to 5hmC accumulation during differentiation at promoters were mostly driven by Group I genes, which associated with glial fate specification and calcium ion homeostasis (**Figure 5B, C**). Coversely, Group II genes were mainly responsible for gene bodies acquiring 5hmC and were linked to proliferative capacity, cell cycle progression, mitochondrial organization, and neural differentiation (**Figure 5B, C**). Enhancers gaining 5hmC also mapped to Group II genes, but were primarily associated with stemness maintenance (**Figure 5B, C**). These findings indicate that TET2-dependent 5hmC deposition segregates into two complementary programs: strong, promoter-associated 5hmC supporting differentiation and calcium signalling, and partially TET2-dependent gene body/enhancer 5hmC maintaining proliferative and metabolic potential, thereby balancing stemness and differentiation.

To further study transcriptional changes associated with 5hmC gain, we analyzed the expression levels of specific genes within the identified biological processes using RNA-seq datasets from wild-type and *Tet2*-deficient NSCs during differentiation. Most genes with altered 5hmC deposition at promoters, gene bodies, or enhancers in *Tet2*-deficient NSCs also displayed expression changes compared to wild-type NSCs (**Figure 6A**). Notably, the number of differentially expressed genes gaining 5hmC at promoters or gene bodies exceeded those gaining 5hmC at enhancers, indicating that promoter- and gene body-associated 5hmC is more tightly linked to transcriptional regulation (**Figure 6A**). In *Tet2*-deficient NSCs, key genes such as *Fgf2,* a regulator of lineage commitment, and *S100b,* associated with astrocytic differentiation, both failed to acquire 5hmC at their promoters (**Figure 6B**). Similarly, the calcium-related genes *Micu2* and *Atp1a2*, which control mitochondrial Ca²⁺ uptake and ionic homeostasis, did not acquire the expected promoter-associated 5hmC (**Figure 6B**). *Foxc1,* transcription factor associated with stemness, did not gain 5hmC within its gene body, confirming broad TET2 dependence at critical regulatory regions. Validation by qPCR in *Tet2*-deficient NSCs differentiated for 3 DIV showed that *Fgf2*, *S100b*, and *Micu2*, which lose 5hmC in their respective regulatory regions, all exhibited significantly reduced expression compared with controls (**Figure 6C**). These results demonstrate that TET2-dependent 5hmC deposition at promoters and gene bodies directly supports the proper acquisition of differentiation-associated transcriptional states.

**Figure 6.**
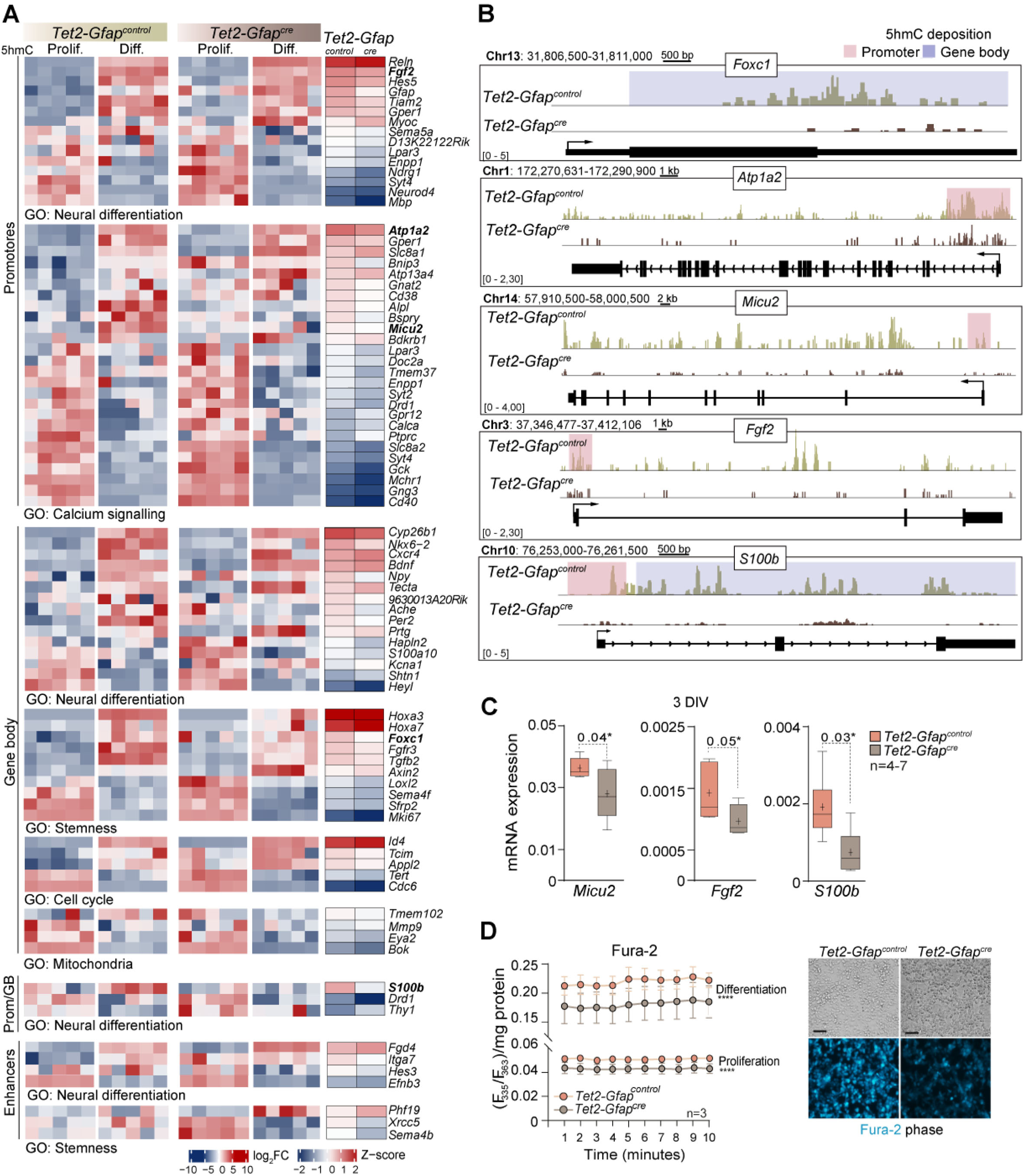
*Tet2* deficiency-induced hypohydroxymethylation leads to transcriptional and functional alterations in adult NSCs. **A)** Heatmap showing Z-score-scaled expression of genes that gain 5hmC during differentiation of *Tet2-Gfap^control^* NSCs, stratified by the genomic location of the 5hmC gain (promoters, gene bodies, or enhancers) and by GO category. Log FoldChanges between proliferating (Prolif.) and 3 DIV-differentiated (Diff.) NSCs are shown for both genotypes. Only genes with an absolute Log FoldChange difference >0.5 between genotypes are displayed. *Phg19, Xrcc5, Sema4b, s100b, Drd1* and *Thy1* are displayed on a different Log FoldChange scale (-2 to 2). **B)** IGV views of 5hmC signal tracks for genes whose 5hmC deposition and transcription are altered in 3 DIV-differentiated (Diff.) *Tet2-Gfap^control^* and *Tet2-Gfap^cre^* NSCs. Genomic positions based on the Genome Reference Consortium Mouse Build 38 (GRCm38/mm10) are also indicated. **C)** qPCR validation of selected genes that gain 5hmC and are upregulated during adult NSCs differentiation but fail to do it in the absence of *Tet2*. NSCs from both genotypes after 3 DIV of differentiation were analyzed. **D)** Intracellular calcium levels measured using Fura-2 ratiometric imaging (F₃₄₀/F₃₈₀) in *Tet2*-*Gfap^control^* and *Tet2*-*Gfap^cre^* NSCs after 3 DIV of differentiation. The graph shows average intracellular calcium levels over 10 minutes, normalized to total protein (mg) (left panel). Contrast images (upper panel) and fluorescence images (bottom panel) of proliferating *Tet2*-*Gfap^control^* and *Tet2*-*Gfap^cre^* NSCs loaded with Fura-2 (blue) (right panel). *Gapdh* was used to normalized gene expression data. DIV: days *in vitro*. In box and whiskers plots, the mean is marked with a (+) and whiskers represent the maximum and minimum values. In D, mean and s.e.m. are shown. Significance was evaluated using linear regression and unpaired one-tailed t-test. P-values and number of samples (n) are indicated. *: P<0.05; ****: P<0.0001, n.s.: not significant. Scale bar in D: 60 μm.

It is known that intracellular calcium regulates cytoskeletal remodeling and astrocyte maturation (Agulhon et al., 2008; Zhou et al., 2019), and calcium-related genes are highly controlled by TET2 *via* promoter-associated 5hmC deposition and transcriptional regulation. To assess one of the functional consequences of these transcriptional and epigenetic defects, we measured basal intracellular calcium levels using Fura-2 ratiometric imaging (Martinez-Zaguilan et al., 1996). In wild-type NSCs, basal calcium levels increased markedly during differentiation, reflecting normal activation of calcium-regulated pathways (**Figure 6D**). In contrast, *Tet2*-deficient NSCs exhibited significantly reduced calcium levels both under proliferative and differentiating conditions, and failed to reach the normal differentiation-associated increase (**Figure 6D**), consistent with the impaired expression of calcium-regulating genes. This deficit provides a mechanistic link between TET2-dependent 5hmC deposition, transcriptional regulation, and the previously observed defects in astrocytic maturation and function in *Tet2*-deficient NSCs.

Together, these results demonstrate that TET2-dependent, locus-specific 5hmC deposition orchestrates transcriptional programs and cellular pathways, such as calcium signalling, as well as genes promoting differentiation and self-renewal, that are essential for the maintenance, metabolic competence, and differentiation of adult NSCs, thereby balancing stemness and lineage progression through coordinated epigenetic and transcriptional regulation.

## Discussion

Adult neurogenesis is a tightly regulated process that relies on the precise coordination of activation, proliferation, and differentiation of neural stem cells (NSCs) within specialized brain niches. Emerging evidence has implicated Ten-eleven translocation (TET) enzymes in the epigenetic regulation of these processes (Gu et al., 2011; Li et al., 2017; Montalbán-Loro et al., 2019; Wu & Zhang, 2017). Our work identifies TET2 as a central regulator of adult NSC homeostasis in the ventricular-subventricular zone (V-SVZ). Using a conditional *Tet2* knockout mouse model, we show that loss of TET2 selectively depletes quiescent NSCs, likely by accelerating their transition into a primed or activated state, without broadly impairing progenitor proliferation. This observation supports a model in which TET2 restrains premature NSC activation, maintaining long-term stem cell pool while preventing overexpansion of proliferating progenitors, consistent with previous findings in other stem cell systems (An et al., 2015; Moran-Crusio et al., 2011; Yuita et al., 2023). Functionally, *Tet2-*deficient NSCs exhibit impaired terminal astrocytic differentiation, resulting in stalled maturation and reduced neurogenesis in the olfactory bulb, underscoring the essential role of TET2 in adult neurogenic output. *In vitro* astrocytic differentiation assays of *Tet2*-deficient adult NSCs corroborated the *in vivo* phenotypes.

Transcriptomic profiling revealed that *Tet2*-deficient NSCs showed defective induction of lineage-specific transcriptional programs, with more pronounced alterations during differentiation than proliferation. Dysregulated genes included those involved in differentiation, metabolic processes, and intracellular calcium signaling, highlighting TET2’s role in coordinating transcriptional programs that integrate stemness, lineage commitment, and functional output. Mechanistically, these transcriptional defects are linked to locus-specific alterations of 5hmC which is known to be enriched in quiescent, non-proliferating, and differentiated cells (López-Moyado et al., 2024).

Genome-wide mapping demonstrated low basal 5hmC levels in proliferating NSCs, consistent with replication-associated dilution and low 5mC substrate at highly expressed promoters in self-renewing cells (Prikrylova et al., 2019). Upon differentiation, wild-type NSCs exhibit robust 5hmC gains at promoters, gene bodies, and enhancers, coinciding with increased TET2 expression. Importantly, promoter- and gene body-associated 5hmC gains were strongly linked to transcriptional regulation. Integration of 5hmC and RNA-seq data revealed that *Tet2*-deficient NSCs fail to achieve comparable hydroxymethylation both in proliferating and differentiating states, delineating two complementary programs: promoter-associated 5hmC gains that are strictly TET2-dependent regulating glial fate specification and calcium homeostasis, and gene body- and enhancer-associated 5hmC gains that are partially TET2-dependent and associated to stemness, proliferation, and metabolic competence. This differential pattern of 5hmC deposition indicates that TET2 coordinates distinct epigenetic programs to balance stemness and lineage commitment. Notably, TET3-deficient NSCs in the SVZ do not exhibit comparable reductions in 5hmC (Montalbán-Loro et al., 2019), reinforcing the notion that TET2 is the predominant DNA demethylase operating in this niche. In contrast, studies in the SGZ have reported increased 5hmC levels during differentiation, yet *Tet2* loss does not result in a global decrease in 5hmC (X. Li et al., 2017), underscoring a niche-specific functional specialization among TET family members.

Promoter- and gene body-associated 5hmC gains were tightly linked to transcriptional regulation. Key genes such as *Fgf2* and *S100b* failed to acquire promoter 5hmC in *Tet2*-deficient NSCs, resulting in reduced expression and impaired differentiation. Similarly, the stemness-associated transcription factor *Foxc1* showed defective 5hmC acquisition at gene bodies, correlating with transcriptional deficits. A particularly novel finding is the connection between TET2-dependent 5hmC deposition and intracellular calcium signaling. Promoter-associated 5hmC in calcium-regulatory genes ensures proper basal and differentiation-associated calcium levels. In our model, the reduction of basal intracellular calcium in *Tet2*-deficient NSCs could impair their ability to execute proper differentiation programs, contributing to stalled astrocytic maturation and defective neurogenesis. This proves a mechanistic link between TET2-dependent epigenetic remodeling, transcriptional regulation, and impaired astrocytic maturation. More broadly, our findings highlight a conserved role for TET2 in integrating epigenetic regulation with intracellular signaling, consistent with observations in other stem cell systems (Lin et al., 2024; Zhang et al., 2022). Indeed, low basal calcium is associated with maintenance of HSCs and epidermal stem cells, whereas calcium increases drive differentiation (Luchsinger et al., 2019; Umemoto et al., 2018). Parallels can be drawn with cerebellar Purkinje cell differentiation, where sustained TET activity and progressive 5hmC accumulation are required for the acquisition of a mature state.

Our data also reinforce the concept that TET2 integrates catalytic and potentially non-catalytic functions to orchestrate NSC behavior. While our findings emphasize promoter-specific hydroxymethylation as a driver of transcriptional activation, partial 5hmC deposition in *Tet2-*deficient NSCs suggests compensatory activity from other TET family members or additional epigenetic mechanisms emphasizing the robustness of NSC regulatory networks (An et al., 2015; Cimmino et al., 2015; Dawlaty et al., 2013; Yuita et al., 2023). However, TET3 deficiency in SVZ NSCs does not cause detectable 5hmC reduction (Montalbán-Loro et al., 2019), and TET1 is very lowly expressed in adult NSCs suggesting that TET2 may be the principal DNA demethylase in this niche. Additionally, TET2 may modulate transcriptional programs through interactions with chromatin regulators, analogous to roles described for TET1 and TET3 in other contexts, further fine-tuning differentiation and stemness (Chen et al., 2013; Chrysanthou et al., 2022; Joshi et al., 2022).

Overall, our results position TET2 as a central integrator of epigenetic regulation and intracellular signaling in adult NSCs. By coupling promoter- and gene body-specific 5hmC deposition with transcriptional control of differentiation genes, TET2 synchronizes lineage progression with functional signaling pathways. This integration ensures that NSCs maintain stemness while executing proper differentiation programs, linking epigenetic remodeling directly to physiological outcomes in astrocytic maturation and neurogenesis. Finally, our study provides a framework to explore TET2’s broader impact on NSC plasticity, aging, and disease. Futures studies will be needed to explore whether TET2 dysfunction extends to reactive astrocyte states or age-related decline in neurogenesis. Moreover, dissecting the interplay of catalytic and non-catalytic TET2 functions, as well as its cooperation with calcium-regulated transcription factors or chromatin remodelers, will be critical to harnessing TET2 pathways for therapeutic modulation of neurological disorders linked to impaired neurogenesis.

## Resource Availability

### Lead Contact

Further information and requests for resources and reagents should be directed to and will be fulfilled by the lead contact, Sacri R. Ferron

### Materials Availability

This study did not generate new unique reagents.

### Data and Code Availability

#### Data

Raw sequencing data generated in this study have been deposited in the Gene Expression Omnibus (GEO) under the following accession number: GSE312705. Normalized signal tracks are provided as processed information for CMS-IP. For RNA-seq raw counts are provided as processed information. Accession numbers are listed in the Key Resources Table. All other data supporting the findings of this study are available within the main text and its Supplementary Information.

#### Code

This study did not generate original code. All software and computational tools used for data processing and analysis are cited in the Methods Details section and summarized in Supplementary Table S5. Non-default parameters used for RNA-seq and CMS-IP-seq analyses are specified in the respective analysis subsections.

#### Other

All materials and reagents used in this study are listed in the Key Resources Table. Any additional information required to reanalyze the data reported in this paper is available from the lead contact upon reasonable request.

## Limitations of the study

Although our results identify TET2 as a major regulator of locus-specific 5hmC remodeling in adult SVZ NSCs, some limitations remain. While TET3 does not appear to regulate 5hmC deposition in NSCs, we cannot exclude a contribution from TET1, which despite its low expression may still influence hydroxymethylation at selected promoters, enhancers, or gene bodies. Thus, additional TET-dependent mechanisms may underlie part of the epigenetic and transcriptional changes observed. Moreover, our study primarily addresses the catalytic role of TET2, without assessing its non-catalytic functions, which may also participate in transcriptional regulation. Future work using catalytically inactive mutants and combined TET perturbations will be required to disentangle these activities and define the division of labor among TET enzymes in adult NSCs.

## Acknowledgements

We would like to thank Dr. López-Moyado for technical support and discussion of the data and the Servicio Central de Soporte a la Investigación Experimental (SCSIE-UVEG) for their assistance. This work was supported by grants from Ministerio de Ciencia e Innovación/AEI (PID2019-110045GB-I00, PID2022-142734OB-I00 and EUR2023-143479), Generalitat Valenciana (CIAICO/2024/287), and Instituto de Salud Carlos III (CIBERNED CB06/05/0086) to SRF. LLC (PRE2020-094137) were funded by the Spanish Formación de Personal Investigador (FPI) fellowship program EJV was funded by the Spanish Formación de Profesorado Universitario (FPU) fellowship program (FPU20/00795). Open Access funding was provided by the Ministerio de Ciencia e Innovación. LJAV was supported by a Powell-Bundle Fellowship of the Jacobs School of Engineering, University of California, San Diego. AR acknowledges funding from R35 grant CA210043 and U01 grant 180152 from the National Institutes of Health, USA.

## Author Contribution

Conceptualization: SRF; Methodology: SRF, LLC, EJV, JP, AR; Software: JD, LAV; Validation: LLC; Formal analysis: SRF, LLC, EJV, JP, LAV; Investigation: LLC, EJV; Resources: SRF, AR; Data curation: SRF, AR, LLC, EJV, JP, LAV. Writing original draft: SRF; Writing, review and editing SRF, AR, LLC, EJV, JP, LAV; Supervision: SRF; AR.

## Declaration of generative AI and AI-assisted technologies in the writing process

During the preparation of this work, the authors used ChatGPT in order to improve the language and ensure grammatical accuracy. After using this tool, the authors reviewed and edited the content as needed and take full responsibility for the content of the publication.

## STAR Methods

### EXPERIMENTAL MODEL AND SUBJECT DETAILS

#### Mouse models and in vivo manipulations

The experiments were conducted in 2-to 4-months-old mice. Both GFAP-cre (6.Cg-Tg(Gfap-cre)73.12Mvs/J) and *Tet2^loxp/loxp^* (B6; 129S-Tet2tm1.1laai/J) mice were obtained from Jackson Laboratory and maintained on a C57BL/6J background. Gfap-Cre mice were generated using a 15 kb mouse *Gfap* promoter cassette containing all introns, promoter regulatory elements, exons, and 2 kb of 3′ and 2.5 kb of 5′ flanking regions of the mouse *Gfap* gene (Garcia et al., 2004) . *Gfap* expression is prevented by the removal of a small region in exon 1. *Tet2^loxp/loxp^* mice possess loxP sites flanking exon 3 (Johnson et al., 1995) . Expression of *Cre-recombinase* results in the excision of these regions. To generate specific deletion of *Tet2* in GFAP positive cells, heterozygous Gfap-Cre transgenic animals were bred to *Tet2^loxp/loxp^*. Littermates lacking GFAP-cre were used as control mice. The Jackson laboratory reports that the Gfap-Cre line have *Cre* expression in the male germline (J. Zhang et al., 2013) . To avoid this problem, Gfap-Cre females and *Tet2*^loxp/loxp^ males were used to generate the experimental animals. Animals were genotyped by PCR analysis of DNA, extracted from mouse ear-punch tissue. Mice were maintained in a 12h light/dark cycle with free access to food and water and according to the Animal Care and Ethics committee of the University of Valencia.

### METHOD DETAILS

#### Neurosphere cultures and differentiation assays

Adult 2-to 4-months-old mice were killed by cervical dislocation. To initiate each independent culture, the brains of two different animals were dissected out and the regions containing the V-SVZ from each hemisphere were isolated and washed in Earle’s balanced salt solution (EBSS; Gibco). Tissues were transferred to EBSS containing 12 U/mL papain (Worthington DBA), 0.2 mg/ ml L-cystein (Sigma-Aldrich), 0.2mg/mL EDTA (Sigma-Aldrich), and incubated for 20 min at 37 °C. Tissue was then rinsed in Dulbecco’s modified Eagle’s medium (DMEM)/F12 medium (1:1 v/v; Gibco) and carefully triturated with a fire-polished Pasteur pipette to a single-cell suspension. Isolated cells were collected by centrifugation, resuspended and cultured in NSCs medium: DMEM/F12 medium containing 2 mM L-glutamine (Gibco), 0.6% D-glucose (Sigma-Aldrich), 0.1% sodium bicarbonate (Sigma-Aldrich), 5mM HEPES (Biowest), 9.6 g/mL putrescine (Sigma-Aldrich), 6.3 ng/mL progesterone (Sigma-Aldrich), 5.2 ng/mL sodium selenite (Sigma-Aldrich), 0.025 mg/mL insulin (Sigma-Aldrich), 0.1 mg/mL apo-transferrin (Sigma-Aldrich), 4 mg/mL bovine serum albumin (BSA, Sigma-Aldrich), 2 μg/mL heparin (sodium salt, grade II; Sigma-Aldrich), and supplemented with 20 ng/mL epidermal growth factor (EGF; Gibco) and 10 ng/mL fibroblast growth factor (FGF; Sigma-Aldrich) (Jiménez-Villalba et al., 2024) . Neurospheres were allowed to develop for 6-10 days in a 95% air-5% CO2 humidified atmosphere at 37 °C. For culture expansion, neurospheres were collected, disaggregated using Accutase® (Sigma-Aldrich) and plated at a relatively high density (10,000 cell/cm^2^) in NSCs medium. For self-renewal assays, neurospheres were treated with Accutase® for 10 min, mechanically dissociated to a single-cell suspension and replated at low density (5 cells/μL) in growth medium. Neurospheres were allowed to develop for 5 days in a 95% air-5% CO2 humidified atmosphere at 37 °C. For cell growth assessment, a fraction of the culture at any given passage point, consisting of 250,000 viable cells, was plated and the number of cells generated was determined at the time of the next passage. To generate the accumulated cell growth curves, the ratio of cell production at each subculturing step was multiplied by the number of cells at the previous point of the curve. This procedure was repeated for each passage. For bulk differentiation assays on secondary neurospheres, 50,000 cell/cm^2^ were seeded in Matrigel® (Corning) -coated coverslips and incubated 2 days in culture medium with FGF but without EGF. Medium was then changed with fresh medium without FGF but supplemented with 2% fetal bovine serum (FBS, Biowest) for 5 more days. For primary astrocytes cultures, a homogeneous single-cell suspension of NSCs isolated from the adult V-SVZ, was seeded in astrocyte medium (Control medium + 10% FBS; 0.22 μM filtered before used) at a density of 50,000 cell/cm^2^ cells on Matrigel-coated plates and incubated at 37°C and 5% CO2 for 18 days.

#### Immunohistochemistry and immunocytochemistry analysis

BrdU administration regimes have been previously detailed (Ferron et al., 2007). Briefly, mice were injected intraperitoneally with 50 mg of BrdU (Sigma-Aldrich) per kg of body weight every 2 hours for 12 consecutive hours (7 injections in total) and euthanized 28 days later for label-retaining cells experiments (LRC). For short-retaining experiments (short-BrdU), mice received a single BrdU injection and were euthanized 1h later. Mice were deeply anesthetized and transcardially perfused with 4% paraformaldehyde (PFA, Sigma-Aldrich) in 0.1M phosphate buffer pH 7.4 (PBS). Brains were dissected out, vibratome-sectioned at 40 μm and serially collected. For whole-mount V-SVZ preparations processing, animals were usually non-perfused and dissection was performed with fresh tissue and fixed by immersion in 4% PFA with 0.5% Triton-X-100 at 4°C overnight. Next day, tissue was rinsed three times in 0.5% Triton-X-100 PBS before the immunohistochemical analysis. For BrdU detection, sections were pre-incubated in 2 N HCl for 20 min at 37 °C and neutralized in 0.1M sodium borate (pH 8.5). Tissue sections were washed in PBS and blocked at room temperature (RT) for 1h in PBS with 0.1% Triton X-100 supplemented with 10% FBS and then incubated overnight at 4 °C with primary Detections were performed with fluorescent secondary antibodies for 1h at RT. Nuclei were counterstained with 1 μg/ml of DAPI (Sigma-Aldrich) and the stained samples were mounted with Fluorsave (Millipore). Due to its thickness, V-SVZ whole-mounts were washed three times in PBS cointaining 2% Triton-X-100 for 15 min each and blocked for 2 hours in blocking buffer with 2% Triton-X-100. Primary antibody incubation was done during 48 hours at 4°C prepared in the same blocking buffer. Secondary antibody incubation was done during 2 hours at RT in the same blocking buffer. Images were captured and analyzed with an Olympus FV10i confocal microscope (Olympus). Cell populations were manually counted using the Olympus and ImageJ/Fiji software. Between 3 and 4 coronal sections of the lateral ventricles of each biological replicate were analyzed in immunohystological population studies. For LRC-BrdU neurogenesis studies, and at least 8-10 coronal section in the V-SVZ and in the medial OB were analyzed for neurogenesis studies. For immunocytochemistry, cultures were fixed with 2% PFA 0.1M PBS for 15 min and performed immunocytochemistry as described (Belenguer et al., 2016). For 5hmC detection, sections were pre-incubated in 2 N HCl for 10 min at 37 °C and neutralized in 0.1M sodium borate (pH 8.5). DAPI (1 μg/mL) was used to counterstain DNA. Images were obtained using a FluoView FV10i (Olympus) confocal laser-scanning microscope and were analyzed using FIJI software. Astrocytic cell size (area, μm^2^) was quantified in 7 DIV-differentiated cultures using ImageJ, analyzing at least 30 random fields per culture.

The mean fluorescence intensity (F.I.) of nuclear marker 5hmC was measured in neurospheres for both *Tet2-Gfap^control^* and *Tet2-Gfap^cre^* genotypes after 2 days of differentiation. Thirty captures were analyzed from each animal. Laser setting were first established on control samples and kept throughput the whole experiment. The F.I. was measured on a scale of 0–255 arbitrary units (a.u.). The F.I. for every counted cell was corrected by its particular background value, measured for every counted section, as follows: [((100*cell intensitynx–backgroundx)/(255))*((255 – backgroundx)/100)]. F.I. is represented as frequency histograms that represent the frequency in percentage of cells regarding the intensity of fluorescence in arbitrary units.

#### Flow cytometry analysis

Cell analysis by flow cytometry was done following previously established protocols (Belenguer et al., 2021). The V-SVZ of both brain hemispheres from each mouse were minced and enzymatically digested using the Neural Tissue Dissociation kit (T) in a GentleMACS Octo Dissociator with heaters (Miltenyi). Digested pieces were mechanically dissociated by pipetting up and down 20 times through a plastic Pasteur pipette. Cell suspension was filtered through a 40 μm nylon filter. Cells were pelleted (300*g*, 10 min), resuspended in 100 mL blocking buffer (HBSS without calcium and magnesium (Gibco), 10 mM HEPES, 2 mM EDTA, 0.1% glucose, 0.5% BSA) and incubated with specific primary antibodies at 4 °C for 30 min. Then, labelled samples were washed with 1 mL of blocking buffer and centrifuged at 300*g* during 10 min. Finally, samples were resuspended in 0.5 mL of blocking buffer and analyzed with a LSR-Fortessa cytometer (Becton Dickinson) with 350, 488, 561 and 640 nm lasers. Dead cells were excluded by staining with 0.1 mg/mL DAPI prior to analysis. For apoptosis evaluation, flow cytometry analysis was performed. Apoptotic cells were identified as a distinct population characterized by reduced forward scatter (FSC), indicative of decreased cell size, and lower side scatter (SSC), reflecting diminished internal complexity.

#### Intracellular Ca^2+^ determination

To estimate the intracellular Ca^2+^ levels in NSCs, the fluorescence probe Fura-2 (acetoxymethyl (AM)-derivative; Invitrogen) was used. NSCs were seed on p96 Matrigel®-treated wells under proliferation conditions in complete medium and under differentiation conditions following the protocol explained above. Four-days proliferating NSCs and 3 days *in vitro* (DIV) under differentiation condition NSCs were added Fura-2 (2 μM in DMSO) to the medium and were incubated for 40 min at 37 °C. Then, cells were washed and further incubated with standard buffer (140 mM NaCl, 2.5 mM KCl, 15 mM Tris-HCl, 5 mM d-glucose, 1.2 mM Na2HPO4, 1 mM MgSO4 and 1 mM CaCl2, pH 7.4) for 30 min at 37 °C. Finally, the standard buffer was removed and experimental buffer (140 mM NaCl, 2.5 mM KCl, 15 mM Tris-HCl, d-glucose, 1.2 mM Na2HPO4, and 2 mM CaCl2, pH 7.4) was added. Emissions at 510 nm, after excitations at 335 and 363 nm, respectively, were recorded in a BioTek Synergy H1 (Agilent) spectrofluorometer at 37 °C. Intracellular calcium levels were estimated by representing the ratio of fluorescence emitted at 510 nm obtained after excitation at 335 nm divided by that at 363 nm (F335/F363). To account for differences in cell density across cultures, fluorescence values were normalized to total protein content (mg protein). To do so, immediately after imaging, cells from each well were lysed with cold lysis buffer (150 mM NaCl, 0.5% sodium deoxycholate, 50 mM Tris-HCl pH 8.0, 1% Tx-100 and 1% SDS) and protein concentration was determined using a BCA protein assay kit (Thermo Fisher). Background subtraction was accomplished from emission values obtained from cells treated with the vehicle (DMSO). At least 6 wells were recorded *per* biological replicate and condition.

#### Expression studies

Total RNA from tissues was prepared with TRIzol reagent (Sigma) per manufacturer’s instructions followed by DNAse (Thermo Scientific) treatment. Briefly, 1 mL of reagent was added per 5-10x10^6^ cells for lysis during 20 min at RT. Then, 100 μL of chloroform was added to samples and incubated at RT for 10 min and centrifuged at 12,000*g* centrifugation at 4°C for 10 min. For RNA precipitation, aqueous phase was mixed with 500 μL of isopropanol and incubated for 5 min. Samples were centrifuged 8 min at 12,000*g* at 4 °C. RNA pellet was washed in 1 mL of 75% ethanol, vortexed and centrifuged 5 min at 7500*g* at 4 °C. Then, RNA pellet was resuspended in RNase-free water and stored at -80°C until use. RNAs from cells were extracted with NZY Total RNA Isolation Kit (NZYtech) including DNase treatment, following the manufacturer’s guidelines. For quantitative PCR, 0.5-1 μg of total RNA was reverse transcribed using the PrimeScript RT reagent kit (Takara) in a final volume of 20μL using random hexamers, following standard procedures. Gene expression analysis was assessed by real-time PCR in a Step One Plus real-time PCR (Applied Biosystem) machine, using 4-10 ng of cDNA contained in 1 μL and specific SYBR-green primers or TaqMan^TM^ probes (Applied Biosystems) for each gene. For TaqMan assays, TaqMan^TM^ Fast Advanced Master Mix (Applied Biosystems) was employed in a 10 μL of final volume. For SYBR-green, reaction included TB Green Premix ExTaq™ (Takara), 0.2 μM of each primer and ROX reference dye in a 10 μL of final volume. All qPCR reactions were performed in a Step One Plus cycler using a standard 40 cycles program with annealing and extension steps at 60°C. Relative expression levels were calculated using the 2–ΔCt method to calculate the relative gene expression compared to the endogenous expression of *Gapdh*.

#### RNA-seq and CMS-IP

Proliferative neurospheres and NSCs differentiated for 3 DIV were harvested for RNA and DNA extraction. RNA Library preparation and high-throughput sequencing were performed by the company Novogene. Five independent cultures *per* condition and genotype were sequenced. Read quality was assessed with FastQC. CMS-IP was performed as previously described (Huang et al., 2012). Briefly, DNA was extracted with DNeasy Blood and Tissue Kit (Qiagen) following the manufacturer’s instructions. Three μg of genomic DNA was sheared using Covaris S2 and was spiked with two 210-base-pair PCR amplicons (C amplicon and 5hmC amplicon) at a ratio of around 1:20,000. DNA was purified with Ampure XP beads (Beckman Coulter) and processed with NEBNext End Repair and dA-Tailing Modules (NEB), followed by a ligation to methylated Illumina Adaptors using NEBNext Quick Ligation Module (NEB). DNA was then treated with sodium bisulfite (MethylCode, Applied Biosystems) for 4 hours, denatured, and immunoprecipitated with anti-CMS serum (input samples were reserved as 1% of total DNA before immunoprecipitation). Samples for immunoprecipitated DNA and input DNA were then purified using Phenol/Chloroform and amplified with index primers using KAPA HiFi HotStart Uracil+ Ready Mix (KAPA Biosystems). The samples after PCR amplification were then purified using Ampure XP Beads. DNA libraries were prepared using NEBNExt Multiplex Oligos for enzymatic Methyl-seq (NEB) and quantified using the TapeStation (Agilent), pooled in equimolar proportions and sequenced as 50-base-pair paired end reads using Illumina Hiseq 2500 (Illumina) to achieve of 25 million reads per sample (n=2-3 independent cultures *per* condition and genotype were sequenced).

The spike-in amplicons were generated using λ phage DNA (Promega) as template and with dNTP mix or 5-Hydroxymethylcytosine dNTP mix (Zymo Research), respectively. Therefore, the cytosines in the C amplicon were unmethylated (used to monitor the bisulfite conversion efficiency), while the cytosines in the 5hmC amplicon were 5-hydroxymethylated (used for normalization of the number of reads to the global content of 5hmC levels in different samples). Reads that mapped to the 5hmC spike-in control in the CMS-IP samples reflected the global content of 5hmC in each sample.

#### RNA-Seq bioinformatics analysis

Initial read quality assessment was performed with FastQC. Expression was quantified at gene level with salmon (Patro et al., 2017) in pseudomapping mode, with automatic library detection (-l A) and sequence bias correction (--seqBias) using the Gencode release M23 as reference (Frankish et al., 2020) . Gene expression quantification was imported into R with package tximeta (Love et al., 2020) and differential expression analysis was performed with DESeq2 (Love et al., 2014) . Experimental batches were corrected with ComBat-ref (X. Zhang, 2025) . Gene ontology (GO) analysis was conducted with R package clusterProfiler (T. Wu et al., 2021) . Tidyverse packages ggplot2, dplyr, tibble and tidyr were used for data wrangling and visualization (Wickham et al., 2019) . Heatmap visualizations were performed with Complex Heatmap package (Z. Gu et al., 2016) and scaled variance stabilized counts.

#### CMS-IP bioinformatics analysis

Initial read quality assessment was performed with FastQC. Sequencing adapters trimmed using Trim Galore (v0.4.2) and low-quality reads were discarded (-q 20). Filtered raw reads were mapped against the *in silico* bisulfite converted mm10 genome with the unmethylated and 5hmC amplicons added as artificial chromosomes with bsmap (Xi & Li, 2009) . PCR duplicates were marked with Samtools (v1.7) (H. Li et al., 2009). Normalization was performed using the 5hmC spike-ins. Specifically, the proportion (Ps) of 5hmC spike-in reads to non-duplicate autosomal reads were determined for each sample (S) as follows:

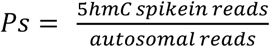

This proportion of 5hmC spike-in reads was used to create a 5hmC normalization ratio for CMS-IP and input samples separately (Ns), using the following formula:

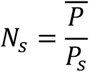

where Ns represents the average proportion of 5hmC reads of all CMS-IP or all input samples. A final 5hmC normalization ratio (Nf) was calculated by correcting the 5hmC normalization ratio of CMS-IP samples (Nc) by the 5hmC normalization ratio for their respective input samples (Ni) as follows:

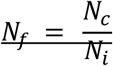

Subsequently, CMS-IP reads were normalized for reads per million multiplied by the final 5hmC normalization ratio (N_f_). Signal tracks were generated with bamCoverage (deepTools suite) (Ramírez et al., 2014) with the flags --scaleFactor N_f_ (previously computed values) and --centerReads. Heatmaps were generated using computeMatrix and plotHeatmap (deepTools suite). GO analyses were performed with clusterProfiler. Differential 5hmC signal analyses were carried out using edgeR (Robinson et al., 2009) with the glmLRT function, and regions with *P value* < 0.05 were considered significant. Promoters were defined as –1Kb, to +500bp from the TSS at the gene level. NSC enhancer annotations was obtained by combining SEdb 3.0 release and Enhancer Atlas 2.0 release (Gao & Qian, 2019; Wang et al., 2022). Enhancers retrieved from Enhancer Atlas were lifted over to mm10 genome, and the resulting genomic coordinates were merged using bedtools merge -d 500. Genes with a TSS located within ±50 kb of an enhancer were considered to be regulated by that enhancer.

### QUANTIFICATION AND STATISTICAL ANALYSIS

All statistical tests were performed using the GraphPad Prism Software, version 9.00 for Windows. Data were first tested for normality using Shapiro-Wilk test. The significance of differences between groups was evaluated using appropriate statistical tests for each comparison. For data that passed normality tests, *t-tests* were used for two group comparison (with Welch’s correction when variances differed), and one-way ANOVA followed by Šídák’s post hoc test was applied for comparisons involving three or more groups. For non-normally distributed data, the Mann–Whitney nonparametric test was used for two-group comparisons. For analysis involving relative values (percentages), data were normalized using arcsine-root transformation. Values of *P* ≤0.05 were considered statistically significant. Data are presented as the mean ± standard error of the mean (s.e.m.). In box-and-whiskers plots, horizontal lines represent Q3, median, and Q1; (+) indicates the mean, and whiskers denote minimum and maximum values. In bar plots, mean and s.e.m are shown. In violin plots, the median (solid line) and first and third quartiles (dashed lines) are indicated. The number of independent cultures or animals (n, dots) and *P* values are shown in the figures: P<0.05 (*); P<0.01 (**); P<0.001 (***); P<0.0001 (****); n.s.; not significant.

## Supplemental data legends

**Supplementary data 1 (Results)**, related to Figures 1-6 and Figures S1-5.

Source data supporting the results shown in the main and supplementary figures.

**Supplementary data 2 (RNA-seq Analysis)**, related to Figures 4- 6 and Figures S4-5.

Differential expression analysis performed using DESeq2. Results are provided in four separate tables corresponding to: (2.1) wild-type differentiated (3 DIV) NSCs vs wild-type proliferating NSCs; (2.2) *Tet2*-deficient differentiated (3 DIV) NSCs vs *Tet2*-deficient proliferating NSCs; (2.3) *Tet2*-deficient proliferating NSCs vs wild-type NSCs proliferating; (2.4) *Tet2*-deficient differentiated (3 DIV) NSCs vs wild-type differentiated (3 DIV) NSCs. Each table includes a detailed description of the results.

**Supplementary data 3 (CMS-IP Analysis),** related to Figures 5-6 and Figure S5. CMS-IP differential signal analysis. Results are provided in six tables, corresponding to promoter-, gene body-, and enhancer-level differential analyses for both wild-type and *Tet2*-deficient NSCs (proliferating vs differentiated, 3 DIV). Each table includes a detailed description of the results. GO results are also presented as a separate tab in this supplement.

**Supplementary data 4 (Gene ontology Analysis),** related to Figures 5-6 and Figure S5. GO results are presented as a separate tab in this supplement. GO term enrichment analysis for promoters, enhancers and gene bodies are analyzed.

## References

1. Agulhon, C., Petravicz, J., McMullen, A. B., Sweger, E. J., Minton, S. K., Taves, S. R., Casper, K. B., Fiacco, T. A. & McCarthy, K. D. (2008). What Is the Role of Astrocyte Calcium in Neurophysiology? Neuron, 59(6), 932–946. 10.1016/j.neuron.2008.09.004

2. An, J., González-Avalos, E., Chawla, A., Jeong, M., López-Moyado, I. F., Li, W., Goodell, M. A., Chavez, L., Ko, M. & Rao, A. (2015). Acute loss of TET function results in aggressive myeloid cancer in mice. Nature Communications, 6(1), 10071. 10.1038/ncomms10071

3. Atlasi, Y. & Stunnenberg, H. G. (2017). The interplay of epigenetic marks during stem cell differentiation and development. Nature Reviews Genetics, 18(11), 643–658. 10.1038/nrg.2017.57

4. Bai, D., Zhang, X., Xiang, H., Guo, Z., Zhu, C. & Yi, C. (2025). Simultaneous single-cell analysis of 5mC and 5hmC with SIMPLE-seq. Nature Biotechnology, 43(1), 85–96. 10.1038/s41587-024-02148-9

5. Belenguer, G., Domingo-Muelas, A., Ferrón, S. R., Morante-Redolat, J. M. & Fariñas, I. (2016). Isolation, culture and analysis of adult subependymal neural stem cells. Differentiation, 91(4–5), 28–41. 10.1016/j.diff.2016.01.005

6. Belenguer, G., Duart-Abadia, P., Domingo-Muelas, A., Morante-Redolat, J. M. & Fariñas, I. (2021). Cell population analysis of the adult murine subependymal neurogenic lineage by flow cytometry. STAR Protocols, 2(2), 100425. 10.1016/j.xpro.2021.100425

7. Bogdanović, O., Smits, A. H., Mustienes, E. de la C., Tena, J. J., Ford, E., Williams, R., Senanayake, U., Schultz, M. D., Hontelez, S., Kruijsbergen, I. van, Rayon, T., Gnerlich, F., Carell, T., Veenstra, G. J. C., Manzanares, M., Sauka-Spengler, T., Ecker, J. R., Vermeulen, M., Gómez-Skarmeta, J. L. & Lister, R. (2016). Active DNA demethylation at enhancers during the vertebrate phylotypic period. Nature Genetics, 48(4), 417–426. 10.1038/ng.3522

8. Cebrian-Silla, A., Nascimento, M. A., Redmond, S. A., Mansky, B., Wu, D., Obernier, K., Rodriguez, R. R., Gonzalez-Granero, S., García-Verdugo, J. M., Lim, D. A. & Álvarez-Buylla, A. (2021). Single-cell analysis of the ventricular-subventricular zone reveals signatures of dorsal and ventral adult neurogenesis. eLife, 10, e67436. 10.7554/elife.67436

9. Chaker, Z., Codega, P. & Doetsch, F. (2016). A mosaic world: puzzles revealed by adult neural stem cell heterogeneity. Wiley Interdisciplinary Reviews: Developmental Biology, 5(6), 640–658. 10.1002/wdev.248

10. Chen, Q., Chen, Y., Bian, C., Fujiki, R. & Yu, X. (2013). TET2 promotes histone O-GlcNAcylation during gene transcription. Nature, 493(7433), 561–564. 10.1038/nature11742

11. Chrysanthou, S., Tang, Q., Lee, J., Taylor, S. J., Zhao, Y., Steidl, U., Zheng, D. & Dawlaty, M. M. (2022). The DNA dioxygenase Tet1 regulates H3K27 modification and embryonic stem cell biology independent of its catalytic activity. Nucleic Acids Research, 50(6), 3169–3189. 10.1093/nar/gkac089

12. Cimmino, L., Dawlaty, M. M., Ndiaye-Lobry, D., Yap, Y. S., Bakogianni, S., Yu, Y., Bhattacharyya, S., Shaknovich, R., Geng, H., Lobry, C., Mullenders, J., King, B., Trimarchi, T., Aranda-Orgilles, B., Liu, C., Shen, S., Verma, A. K., Jaenisch, R. & Aifantis, I. (2015). TET1 is a tumor suppressor of hematopoietic malignancy. Nature Immunology, 16(6), 653–662. 10.1038/ni.3148

13. Cui, X.-L., Nie, J., Ku, J., Dougherty, U., West-Szymanski, D. C., Collin, F., Ellison, C. K., Sieh, L., Ning, Y., Deng, Z., Zhao, C. W. T., Bergamaschi, A., Pekow, J., Wei, J., Beadell, A. V., Zhang, Z., Sharma, G., Talwar, R., Arensdorf, P., … He, C. (2020). A human tissue map of 5-hydroxymethylcytosines exhibits tissue specificity through gene and enhancer modulation. Nature Communications, 11(1), 6161. 10.1038/s41467-020-20001-w

14. Dall’Agnese, A. & Young, R. (2023). Regulatory architecture of cell identity genes and housekeeping genes. Trends in Cell Biology, 33(12), 1010–1013. 10.1016/j.tcb.2023.08.007

15. Dawlaty, M. M., Breiling, A., Le, T., Raddatz, G., Barrasa, M. I., Cheng, A. W., Gao, Q., Powell, B. E., Li, Z., Xu, M., Faull, K. F., Lyko, F. & Jaenisch, R. (2013). Combined Deficiency of Tet1 and Tet2 Causes Epigenetic Abnormalities but Is Compatible with Postnatal Development. Developmental Cell, 24(3), 310–323. 10.1016/j.devcel.2012.12.015

16. Delgado, A. C., Maldonado-Soto, A. R., Silva-Vargas, V., Mizrak, D., Känel, T. von, Tan, K. R., Paul, A., Madar, A., Cuervo, H., Kitajewski, J., Lin, C.-S. & Doetsch, F. (2021). Release of stem cells from quiescence reveals gliogenic domains in the adult mouse brain. Science, 372(6547), 1205–1209. 10.1126/science.abg8467

17. Doetsch, F. (2003). The glial identity of neural stem cells. Nature Neuroscience, 6(11), 1127–1134. 10.1038/nn1144

18. Figueres-Oñate, M., Sánchez-Villalón, M., Sánchez-González, R. & López-Mascaraque, L. (2019). Lineage Tracing and Cell Potential of Postnatal Single Progenitor Cells In Vivo. Stem Cell Reports, 13(4), 700–712. 10.1016/j.stemcr.2019.08.010

19. Frankish, A., Diekhans, M., Jungreis, I., Lagarde, J., Loveland, J. E., Mudge, J. M., Sisu, C., Wright, J. C., Armstrong, J., Barnes, I., Berry, A., Bignell, A., Boix, C., Sala, S. C., Cunningham, F., Domenico, T. D., Donaldson, S., Fiddes, I. T., Girón, C. G., … Flicek, P. (2020). GENCODE 2021. Nucleic Acids Research, 49(D1), D916–D923. 10.1093/nar/gkaa1087

20. Friedman, M. J., Wagner, T., Lee, H., Rosenfeld, M. G. & Oh, S. (2024). Enhancer–promoter specificity in gene transcription: molecular mechanisms and disease associations. Experimental & Molecular Medicine, 56(4), 772–787. 10.1038/s12276-024-01233-y

21. Gao, T. & Qian, J. (2019). EnhancerAtlas 2.0: an updated resource with enhancer annotation in 586 tissue/cell types across nine species. Nucleic Acids Research, 48(D1), D58–D64. 10.1093/nar/gkz980

22. Garcia, A. D. R., Doan, N. B., Imura, T., Bush, T. G. & Sofroniew, M. V. (2004). GFAP-expressing progenitors are the principal source of constitutive neurogenesis in adult mouse forebrain. Nature Neuroscience, 7(11), 1233–1241. 10.1038/nn1340

23. Globisch, D., Münzel, M., Müller, M., Michalakis, S., Wagner, M., Koch, S., Brückl, T., Biel, M. & Carell, T. (2010). Tissue Distribution of 5-Hydroxymethylcytosine and Search for Active Demethylation Intermediates. PLoS ONE, 5(12), e15367. 10.1371/journal.pone.0015367

24. Gu, T.-P., Guo, F., Yang, H., Wu, H.-P., Xu, G.-F., Liu, W., Xie, Z.-G., Shi, L., He, X., Jin, S., Iqbal, K., Shi, Y. G., Deng, Z., Szabó, P. E., Pfeifer, G. P., Li, J. & Xu, G.-L. (2011). The role of Tet3 DNA dioxygenase in epigenetic reprogramming by oocytes. Nature, 477(7366), 606–610. 10.1038/nature10443

25. Gu, Z., Eils, R. & Schlesner, M. (2016). Complex heatmaps reveal patterns and correlations in multidimensional genomic data. Bioinformatics, 32(18), 2847–2849. 10.1093/bioinformatics/btw313

26. Guo, J. U., Su, Y., Zhong, C., Ming, G. & Song, H. (2011). Hydroxylation of 5-Methylcytosine by TET1 Promotes Active DNA Demethylation in the Adult Brain. Cell, 145(3), 423–434. 10.1016/j.cell.2011.03.022

27. He, B., Zhang, C., Zhang, X., Fan, Y., Zeng, H., Liu, J., Meng, H., Bai, D., Peng, J., Zhang, Q., Tao, W. & Yi, C. (2021). Tissue-specific 5-hydroxymethylcytosine landscape of the human genome. Nature Communications, 12(1), 4249. 10.1038/s41467-021-24425-w

28. He, Y.-F., Li, B.-Z., Li, Z., Liu, P., Wang, Y., Tang, Q., Ding, J., Jia, Y., Chen, Z., Li, L., Sun, Y., Li, X., Dai, Q., Song, C.-X., Zhang, K., He, C. & Xu, G.-L. (2011). Tet-Mediated Formation of 5-Carboxylcytosine and Its Excision by TDG in Mammalian DNA. Science, 333(6047), 1303–1307. 10.1126/science.1210944

29. Honig, F. & Murrell, A. (2025). Cell identity and 5-hydroxymethylcytosine. Epigenetics & Chromatin, 18(1), 36. 10.1186/s13072-025-00601-w

30. Huang, Y., Chavez, L., Chang, X., Wang, X., Pastor, W. A., Kang, J., Zepeda-Martínez, J. A., Pape, U. J., Jacobsen, S. E., Peters, B. & Rao, A. (2014). Distinct roles of the methylcytosine oxidases Tet1 and Tet2 in mouse embryonic stem cells. Proceedings of the National Academy of Sciences, 111(4), 1361–1366. 10.1073/pnas.1322921111

31. Huang, Y., Pastor, W. A., Zepeda-Martínez, J. A. & Rao, A. (2012). The anti-CMS technique for genome-wide mapping of 5-hydroxymethylcytosine. Nature Protocols, 7(10), 1897–1908. 10.1038/nprot.2012.103

32. Ito, S., D’Alessio, A. C., Taranova, O. V., Hong, K., Sowers, L. C. & Zhang, Y. (2010). Role of Tet proteins in 5mC to 5hmC conversion, ES-cell self-renewal and inner cell mass specification. Nature, 466(7310), 1129–1133. 10.1038/nature09303

33. Jiménez-Villalba, E., Lázaro-Carot, L., Mateos-Martínez, C. M., Díaz-Moncho, J., Samper-Llavador, D., Igual-López, M., Planells, J. & Ferrón, S. R. (2024). Isolation, Expansion, and Nucleofection of Neural Stem Cells from Adult Murine Subventricular Zone. Journal of Visualized Experiments : JoVE, 208. 10.3791/66651

34. Johnson, W. B., Ruppe, M. D., Rockenstein, E. M., Price, J., Sarthy, V. P., Verderber, L. C. & Mucke, L. (1995). Indicator expression directed by regulatory sequences of the glial fibrillary acidic protein (GFAP) gene: In vivo comparison of distinct GFAP - lacZ transgenes. Glia, 13(3), 174–184. 10.1002/glia.440130304

35. Joshi, K., Liu, S., S.J., P. B. & Zhang, J. (2022). Mechanisms that regulate the activities of TET proteins. Cellular and Molecular Life Sciences, 79(7), 363. 10.1007/s00018-022-04396-x

36. Kempermann, G., Song, H. & Gage, F. H. (2015). Neurogenesis in the Adult Hippocampus. Cold Spring Harbor Perspectives in Biology, 7(9), a018812. 10.1101/cshperspect.a018812

37. Ko, M., Huang, Y., Jankowska, A. M., Pape, U. J., Tahiliani, M., Bandukwala, H. S., An, J., Lamperti, E. D., Koh, K. P., Ganetzky, R., Liu, X. S., Aravind, L., Agarwal, S., Maciejewski, J. P. & Rao, A. (2010). Impaired hydroxylation of 5-methylcytosine in myeloid cancers with mutant TET2. Nature, 468(7325), 839–843. 10.1038/nature09586

38. Kriaucionis, S. & Heintz, N. (2009). The Nuclear DNA Base 5-Hydroxymethylcytosine Is Present in Purkinje Neurons and the Brain. Science, 324(5929), 929–930. 10.1126/science.1169786

39. Li, H., Handsaker, B., Wysoker, A., Fennell, T., Ruan, J., Homer, N., Marth, G., Abecasis, G., Durbin, R. & Subgroup, 1000 Genome Project Data Processing. (2009). The Sequence Alignment/Map format and SAMtools. Bioinformatics, 25(16), 2078–2079. 10.1093/bioinformatics/btp352

40. Li, L., Gao, Y., Wu, Q., Cheng, A. S. L. & Yip, K. Y. (2019). New guidelines for DNA methylome studies regarding 5-hydroxymethylcytosine for understanding transcriptional regulation. Genome Research, 29(4), 543–553. 10.1101/gr.240036.118

41. Li, X., Yao, B., Chen, L., Kang, Y., Li, Y., Cheng, Y., Li, L., Lin, L., Wang, Z., Wang, M., Pan, F., Dai, Q., Zhang, W., Wu, H., Shu, Q., Qin, Z., He, C., Xu, M. & Jin, P. (2017). Ten-eleven translocation 2 interacts with forkhead box O3 and regulates adult neurogenesis. Nature Communications, 8(1), 15903. 10.1038/ncomms15903

42. Lian, H., Li, W.-B. & Jin, W.-L. (2016). The emerging insights into catalytic or non-catalytic roles of TET proteins in tumors and neural development. Oncotarget, 7(39), 64512–64525. 10.18632/oncotarget.11412

43. Lim, D. A. & Alvarez-Buylla, A. (2016). The Adult Ventricular–Subventricular Zone (V-SVZ) and Olfactory Bulb (OB) Neurogenesis. Cold Spring Harbor Perspectives in Biology, 8(5), a018820. 10.1101/cshperspect.a018820

44. Lin, A. E., Bapat, A. C., Xiao, L., Niroula, A., Ye, J., Wong, W. J., Agrawal, M., Farady, C. J., Boettcher, A., Hergott, C. B., McConkey, M., Flores-Bringas, P., Shkolnik, V., Bick, A. G., Milan, D., Natarajan, P., Libby, P., Ellinor, P. T. & Ebert, B. L. (2024). Clonal Hematopoiesis of Indeterminate Potential With Loss of Tet2 Enhances Risk for Atrial Fibrillation Through Nlrp3 Inflammasome Activation. Circulation, 149(18), 1419–1434. 10.1161/circulationaha.123.065597

45. Llorens-Bobadilla, E., Zhao, S., Baser, A., Saiz-Castro, G., Zwadlo, K. & Martin-Villalba, A. (2015). Single-Cell Transcriptomics Reveals a Population of Dormant Neural Stem Cells that Become Activated upon Brain Injury. Cell Stem Cell, 17(3), 329–340. 10.1016/j.stem.2015.07.002

46. López-Moyado, I. F., Ko, M., Hogan, P. G. & Rao, A. (2024). TET Enzymes in the Immune System: From DNA Demethylation to Immunotherapy, Inflammation, and Cancer. Annual Review of Immunology, 42(1), 455–488. 10.1146/annurev-immunol-080223-044610

47. Love, M. I., Huber, W. & Anders, S. (2014). Moderated estimation of fold change and dispersion for RNA-seq data with DESeq2. Genome Biology, 15(12), 550. 10.1186/s13059-014-0550-8

48. Love, M. I., Soneson, C., Hickey, P. F., Johnson, L. K., Pierce, N. T., Shepherd, L., Morgan, M. & Patro, R. (2020). Tximeta: Reference sequence checksums for provenance identification in RNA-seq. PLoS Computational Biology, 16(2), e1007664. 10.1371/journal.pcbi.1007664

49. Luchsinger, L. L., Strikoudis, A., Danzl, N. M., Bush, E. C., Finlayson, M. O., Satwani, P., Sykes, M., Yazawa, M. & Snoeck, H.-W. (2019). Harnessing Hematopoietic Stem Cell Low Intracellular Calcium Improves Their Maintenance In Vitro. Cell Stem Cell, 25(2), 225–240.e7. 10.1016/j.stem.2019.05.002

50. Martinez-Zaguilan, R., Gurule, M. W. & Lynch, R. M. (1996). Simultaneous measurement of intracellular pH and Ca2+ in insulin-secreting cells by spectral imaging microscopy. American Journal of Physiology-Cell Physiology, 270(5), C1438–C1446. 10.1152/ajpcell.1996.270.5.c1438

51. Montalbán-Loro, R., Lozano-Ureña, A., Ito, M., Krueger, C., Reik, W., Ferguson-Smith, A. C. & Ferrón, S. R. (2019). TET3 prevents terminal differentiation of adult NSCs by a non-catalytic action at Snrpn. Nature Communications, 10(1), 1726. 10.1038/s41467-019-09665-1

52. Moran-Crusio, K., Reavie, L., Shih, A., Abdel-Wahab, O., Ndiaye-Lobry, D., Lobry, C., Figueroa, M. E., Vasanthakumar, A., Patel, J., Zhao, X., Perna, F., Pandey, S., Madzo, J., Song, C., Dai, Q., He, C., Ibrahim, S., Beran, M., Zavadil, J., … Levine, R. L. (2011). Tet2 Loss Leads to Increased Hematopoietic Stem Cell Self-Renewal and Myeloid Transformation. Cancer Cell, 20(1), 11–24. 10.1016/j.ccr.2011.06.001

53. Münzel, M., Globisch, D., Brückl, T., Wagner, M., Welzmiller, V., Michalakis, S., Müller, M., Biel, M. & Carell, T. (2010). Quantification of the Sixth DNA Base Hydroxymethylcytosine in the Brain. Angewandte Chemie International Edition, 49(31), 5375–5377. 10.1002/anie.201002033

54. Obernier, K. & Alvarez-Buylla, A. (2019). Neural stem cells: origin, heterogeneity and regulation in the adult mammalian brain. Development, 146(4), dev156059. 10.1242/dev.156059

55. Obernier, K., Cebrian-Silla, A., Thomson, M., Parraguez, J. I., Anderson, R., Guinto, C., Rodriguez, J. R., Garcia-Verdugo, J.-M. & Alvarez-Buylla, A. (2018). Adult Neurogenesis Is Sustained by Symmetric Self-Renewal and Differentiation. Cell Stem Cell, 22(2), 221–234.e8. 10.1016/j.stem.2018.01.003

56. Palczewski, M. B., Kuschman, H. P., Hoffman, B. M., Kathiresan, V., Yang, H., Glynn, S. A., Wilson, D. L., Kool, E. T., Montfort, W. R., Chang, J., Petenkaya, A., Chronis, C., Cundari, T. R., Sappa, S., Islam, K., McVicar, D. W., Fan, Y., Chen, Q., Meerzaman, D., … Thomas, D. D. (2025). Nitric oxide inhibits ten-eleven translocation DNA demethylases to regulate 5mC and 5hmC across the genome. Nature Communications, 16(1), 1732. 10.1038/s41467-025-56928-1

57. Patro, R., Duggal, G., Love, M. I., Irizarry, R. A. & Kingsford, C. (2017). Salmon provides fast and bias-aware quantification of transcript expression. Nature Methods, 14(4), 417–419. 10.1038/nmeth.4197

58. Prikrylova, T., Robertson, J., Ferrucci, F., Konorska, D., Aanes, H., Manaf, A., Zhang, B., Vågbø, C. B., Kuśnierczyk, A., Gilljam, K. M., Løvkvam-Køster, C., Otterlei, M., Dahl, J. A., Enserink, J., Klungland, A. & Robertson, A. B. (2019). 5-hydroxymethylcytosine Marks Mammalian Origins Acting as a Barrier to Replication. Scientific Reports, 9(1), 11065. 10.1038/s41598-019-47528-3

59. Ramírez, F., Dündar, F., Diehl, S., Grüning, B. A. & Manke, T. (2014). deepTools: a flexible platform for exploring deep-sequencing data. Nucleic Acids Research, 42(W1), W187–W191. 10.1093/nar/gku365

60. Rasmussen, K. D., Berest, I., Keβler, S., Nishimura, K., Simón-Carrasco, L., Vassiliou, G. S., Pedersen, M. T., Christensen, J., Zaugg, J. B. & Helin, K. (2019). TET2 binding to enhancers facilitates transcription factor recruitment in hematopoietic cells. Genome Research, 29(4), 564–575. 10.1101/gr.239277.118

61. Robinson, M. D., McCarthy, D. J. & Smyth, G. K. (2009). edgeR: a Bioconductor package for differential expression analysis of digital gene expression data. Bioinformatics, 26(1), 139–140. 10.1093/bioinformatics/btp616

62. Sohn, J., Orosco, L., Guo, F., Chung, S.-H., Bannerman, P., Ko, E. M., Zarbalis, K., Deng, W. & Pleasure, D. (2015). The Subventricular Zone Continues to Generate Corpus Callosum and Rostral Migratory Stream Astroglia in Normal Adult Mice. The Journal of Neuroscience, 35(9), 3756–3763. 10.1523/jneurosci.3454-14.2015

63. Suh, H., Consiglio, A., Ray, J., Sawai, T., D’Amour, K. A. & Gage, F. H. (2007). In Vivo Fate Analysis Reveals the Multipotent and Self-Renewal Capacities of Sox2 + Neural Stem Cells in the Adult Hippocampus. Cell Stem Cell, 1(5), 515–528. 10.1016/j.stem.2007.09.002

64. Sun, W., Zang, L., Shu, Q. & Li, X. (2014). From development to diseases: The role of 5hmC in brain. Genomics, 104(5), 347–351. 10.1016/j.ygeno.2014.08.021

65. Szulwach, K. E., Li, X., Li, Y., Song, C.-X., Wu, H., Dai, Q., Irier, H., Upadhyay, A. K., Gearing, M., Levey, A. I., Vasanthakumar, A., Godley, L. A., Chang, Q., Cheng, X., He, C. & Jin, P. (2011). 5-hmC–mediated epigenetic dynamics during postnatal neurodevelopment and aging. Nature Neuroscience, 14(12), 1607–1616. 10.1038/nn.2959

66. Szwagierczak, A., Bultmann, S., Schmidt, C. S., Spada, F. & Leonhardt, H. (2010). Sensitive enzymatic quantification of 5-hydroxymethylcytosine in genomic DNA. Nucleic Acids Research, 38(19), e181–e181. 10.1093/nar/gkq684

67. Tahiliani, M., Koh, K. P., Shen, Y., Pastor, W. A., Bandukwala, H., Brudno, Y., Agarwal, S., Iyer, L. M., Liu, D. R., Aravind, L. & Rao, A. (2009). Conversion of 5-Methylcytosine to 5-Hydroxymethylcytosine in Mammalian DNA by MLL Partner TET1. Science, 324(5929), 930–935. 10.1126/science.1170116

68. Tsagaratou, A., Äijö, T., Lio, C.-W. J., Yue, X., Huang, Y., Jacobsen, S. E., Lähdesmäki, H. & Rao, A. (2014). Dissecting the dynamic changes of 5-hydroxymethylcytosine in T-cell development and differentiation. Proceedings of the National Academy of Sciences, 111(32), E3306–E3315. 10.1073/pnas.1412327111

69. Tsagaratou, A., González-Avalos, E., Rautio, S., Scott-Browne, J. P., Togher, S., Pastor, W. A., Rothenberg, E. V., Chavez, L., Lähdesmäki, H. & Rao, A. (2017). TET proteins regulate the lineage specification and TCR-mediated expansion of iNKT cells. Nature Immunology, 18(1), 45–53. 10.1038/ni.3630

70. Umemoto, T., Hashimoto, M., Matsumura, T., Nakamura-Ishizu, A. & Suda, T. (2018). Ca2+–mitochondria axis drives cell division in hematopoietic stem cells. Journal of Experimental Medicine, 215(8), 2097–2113. 10.1084/jem.20180421

71. Urbán, N., Blomfield, I. M. & Guillemot, F. (2019). Quiescence of Adult Mammalian Neural Stem Cells: A Highly Regulated Rest. Neuron, 104(5), 834–848. 10.1016/j.neuron.2019.09.026

72. Wagh, K., Stavreva, D. A. & Hager, G. L. (2025). Transcription dynamics and genome organization in the mammalian nucleus: Recent advances. Molecular Cell, 85(2), 208–224. 10.1016/j.molcel.2024.09.022

73. Wang, Y., Song, C., Zhao, J., Zhang, Y., Zhao, X., Feng, C., Zhang, G., Zhu, J., Wang, F., Qian, F., Zhou, L., Zhang, J., Bai, X., Ai, B., Liu, X., Wang, Q. & Li, C. (2022). SEdb 2.0: a comprehensive super-enhancer database of human and mouse. Nucleic Acids Research, 51(D1), D280–D290. 10.1093/nar/gkac968

74. Wen, L., Li, X., Yan, L., Tan, Y., Li, R., Zhao, Y., Wang, Y., Xie, J., Zhang, Y., Song, C., Yu, M., Liu, X., Zhu, P., Li, X., Hou, Y., Guo, H., Wu, X., He, C., Li, R., … Qiao, J. (2014). Whole-genome analysis of 5-hydroxymethylcytosine and 5-methylcytosine at base resolution in the human brain. Genome Biology, 15(3), R49–R49. 10.1186/gb-2014-15-3-r49

75. Wickham, H., Averick, M., Bryan, J., Chang, W., McGowan, L., François, R., Grolemund, G., Hayes, A., Henry, L., Hester, J., Kuhn, M., Pedersen, T., Miller, E., Bache, S., Müller, K., Ooms, J., Robinson, D., Seidel, D., Spinu, V., … Yutani, H. (2019). Welcome to the Tidyverse. Journal of Open Source Software, 4(43), 1686. 10.21105/joss.01686

76. Wu, T., Hu, E., Xu, S., Chen, M., Guo, P., Dai, Z., Feng, T., Zhou, L., Tang, W., Zhan, L., Fu, X., Liu, S., Bo, X. & Yu, G. (2021). clusterProfiler 4.0: A universal enrichment tool for interpreting omics data. The Innovation, 2(3), 100141. 10.1016/j.xinn.2021.100141

77. Wu, X. & Zhang, Y. (2017). TET-mediated active DNA demethylation: mechanism, function and beyond. Nature Reviews Genetics, 18(9), 517–534. 10.1038/nrg.2017.33

78. Xi, Y. & Li, W. (2009). BSMAP: whole genome bisulfite sequence MAPping program. BMC Bioinformatics, 10(1), 232. 10.1186/1471-2105-10-232

79. Xu, Y., Wu, F., Tan, L., Kong, L., Xiong, L., Deng, J., Barbera, A. J., Zheng, L., Zhang, H., Huang, S., Min, J., Nicholson, T., Chen, T., Xu, G., Shi, Y., Zhang, K. & Shi, Y. G. (2011). Genome-wide Regulation of 5hmC, 5mC, and Gene Expression by Tet1 Hydroxylase in Mouse Embryonic Stem Cells. Molecular Cell, 42(4), 451–464. 10.1016/j.molcel.2011.04.005

80. Yuita, H., López-Moyado, I. F., Jeong, H., Cheng, A. X., Scott-Browne, J., An, J., Nakayama, T., Onodera, A., Ko, M. & Rao, A. (2023). Inducible disruption of Tet genes results in myeloid malignancy, readthrough transcription, and a heterochromatin-to-euchromatin switch. Proceedings of the National Academy of Sciences, 120(6), e2214824120. 10.1073/pnas.2214824120

81. Zhang, H., Wang, S., Zhou, Q., Liao, Y., Luo, W., Peng, Z., Ren, R. & Wang, H. (2022). Disturbance of calcium homeostasis and myogenesis caused by TET2 deletion in muscle stem cells. Cell Death Discovery, 8(1), 236. 10.1038/s41420-022-01041-1

82. Zhang, J., Dublin, P., Griemsmann, S., Klein, A., Brehm, R., Bedner, P., Fleischmann, B. K., Steinhäuser, C. & Theis, M. (2013). Germ-Line Recombination Activity of the Widely Used hGFAP-Cre and Nestin-Cre Transgenes. PLoS ONE, 8(12), e82818. 10.1371/journal.pone.0082818

83. Zhang, R.-R., Cui, Q.-Y., Murai, K., Lim, Y. C., Smith, Z. D., Jin, S., Ye, P., Rosa, L., Lee, Y. K., Wu, H.-P., Liu, W., Xu, Z.-M., Yang, L., Ding, Y.-Q., Tang, F., Meissner, A., Ding, C., Shi, Y. & Xu, G.-L. (2013). Tet1 Regulates Adult Hippocampal Neurogenesis and Cognition. Cell Stem Cell, 13(2), 237–245. 10.1016/j.stem.2013.05.006

84. Zhang, X. (2025). Highly effective batch effect correction method for RNA-seq count data. Computational and Structural Biotechnology Journal, 27, 58–64. 10.1016/j.csbj.2024.12.010

85. Zhou, B., Chen, L., Liao, P., Huang, L., Chen, Z., Liao, D., Yang, L., Wang, J., Yu, G., Wang, L., Zhang, J., Zuo, Y., Liu, J. & Jiang, R. (2019). Astroglial dysfunctions drive aberrant synaptogenesis and social behavioral deficits in mice with neonatal exposure to lengthy general anesthesia. PLoS Biology, 17(8), e3000086. 10.1371/journal.pbio.3000086

