## Supplementary Figures and Tables for "TET2-dependent differential 5hmC deposition balances adult neural stem cell activation and differentiation"

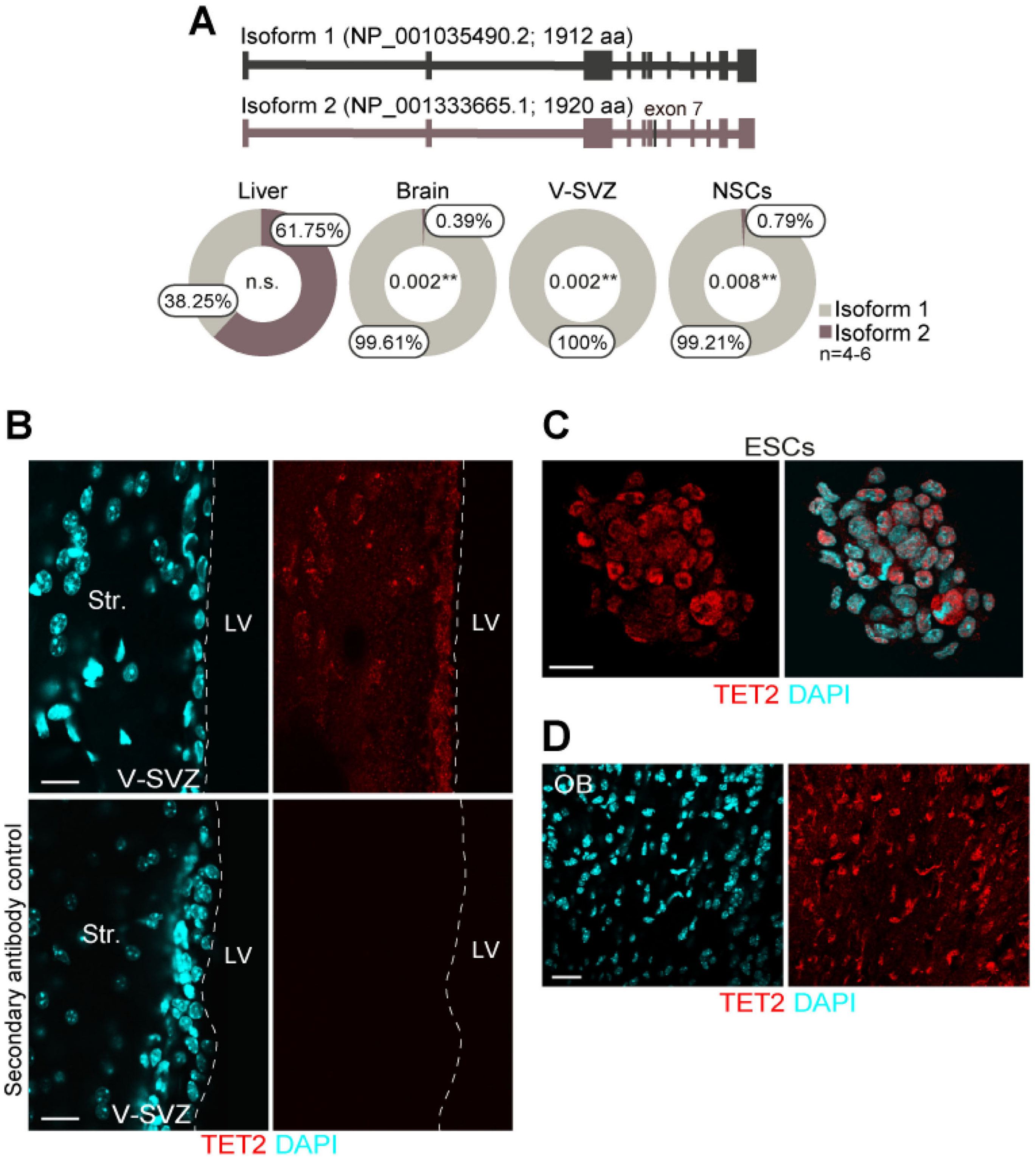


**Supplemental Figure 1. *Tet2* isoform 1 is the predominant transcript expressed in adult NSCs. Related to Figure 1. A)** Schematic representation of the two *Tet2* isoforms identified in mouse (upper panel). Isoform 1 lacks exon 7. The relative abundance of *Tet2* isoforms quantified by qPCR in whole brain, the V-SVZ, and isolated NSCs is shown. Liver tissue was included as a non-neural control (lower panel). *Rps18* served as the normalization control for qPCR analysis. **B)** Immunohistochemistry showing TET2 expression (red) in the V-SVZ of adult wild-type mice (upper panel). No-primary antibody controls (secondary antibody only) show no detectable TET2 signal (lower panel). **C)** TET2 expression (red) in ESCs was used as a positive control. **D)** TET2 expression (red) in the olfactory bulb (OB). DAPI was used to counterstain DNA. V-SVZ: ventricular-subventricular zone; LV: lateral ventricle lumen; Str.: striatal parenchyma; ESC: embryonic stem cell. Statistical significance was assessed using unpaired two-tailed t tests and Mann-Whitney tests. P-values and sample sizes (n) are indicated. **: $P<0.01$; n.s.: not significant. Scale bars in B, C, D: 10 µm.

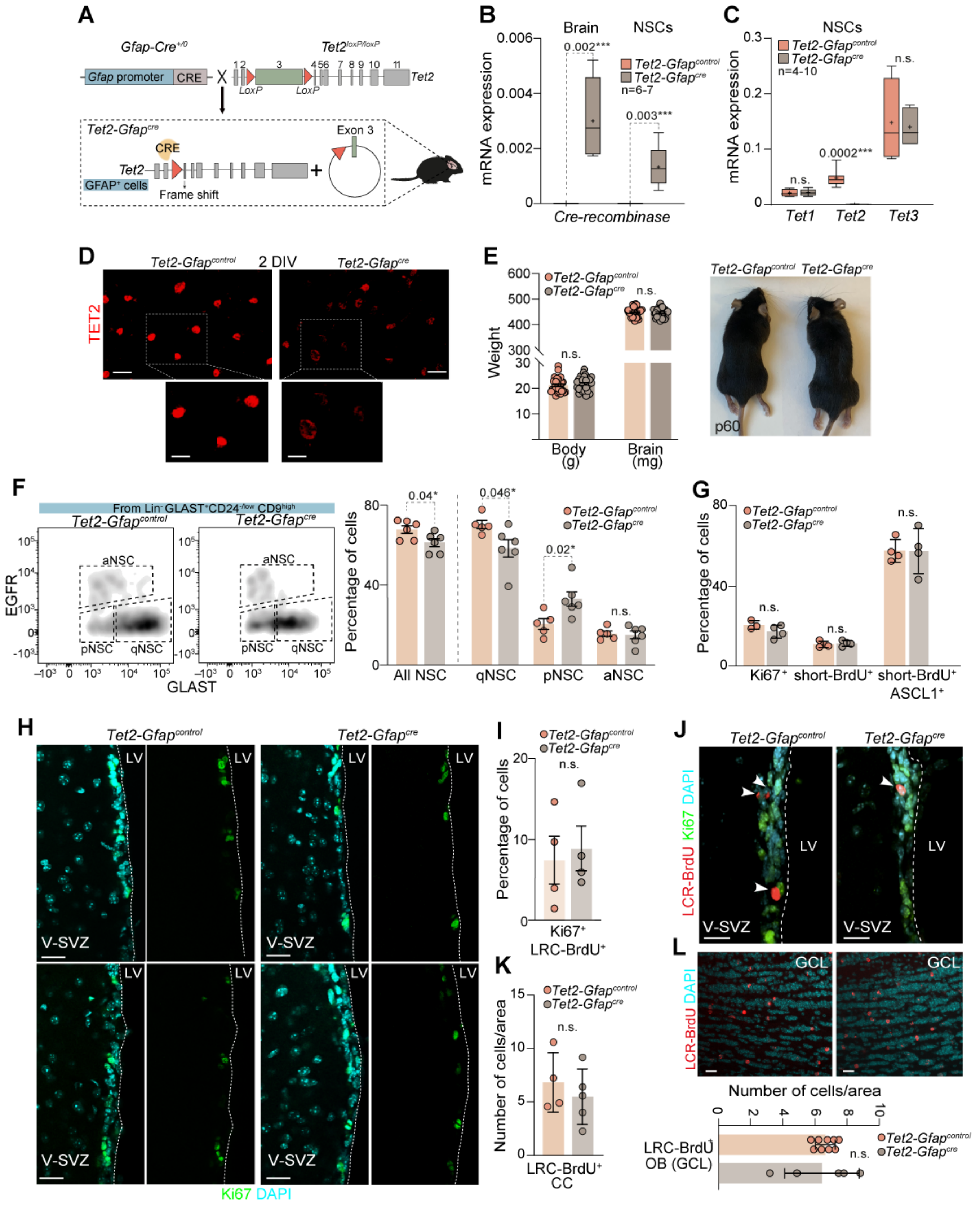


**Supplemental Figure 2. Figure legend provided on the following page. Related to Figure 2.**

**Supplemental Figure 2. Conditional deletion of *Tet2* in GFAP-positive cells using a Cre/LoxP system alters NSCs activation but nor proliferation in the adult V-SVZ. Related to Figure 2. A)** Schematic representation of the mouse model. Mice expressing *Cre recombinase* under the *Gfap* promoter (Gfap-$Cre^{+/0}$) were crossed with mice carrying LoxP sites flanking exon 3 of *Tet2* ($Tet2^{loxP/loxP}$). *Cre*-mediated recombination excises this region, generating a frameshift from exon 3 that disrupts all downstream exons. **B)** qPCR analysis of *Cre* expression in adult brain and NSCs from *Tet2-Gfap*$^{control}$ and *Tet2-Gfap*$^{cre}$ mice. **C)** qPCR analysis of *Tet1, Tet2* and *Tet3* expression in adult NSCs from *Tet2-Gfap*$^{control}$ and *Tet2-Gfap*$^{cre}$ mice. **D)** Immunocytochemistry for TET2 (red) in *Tet2-Gfap*$^{control}$ and *Tet2-Gfap*$^{cre}$ NSCs after 2 DIV under differentiation conditions. **E)** Body weight (grams) and brain weight (milligrams) at postnatal day 60 (p60) in *Tet2-Gfap*$^{control}$ and *Tet2-Gfap*$^{cre}$ mice (left panel). Representative images of *Tet2-Gfap*$^{control}$ and *Tet2-Gfap*$^{cre}$ mice (right panel). **F)** Flow cytometry analysis of GLAST, EGFR and CD9 expression in the V-SVZ of *Tet2-Gfap*$^{control}$ and *Tet2-Gfap*$^{cre}$ mice. NSCs are identified as $GLAST^{+}CD9^{high}$ cells. Subpopulations include quiescent NSCs (qNSCs: $GLAST^{+}/CD9^{high}/EGFR^{low}$), primed NSCs (pNSCs: $GLAST^{low}/CD9^{high}/EGFR^{low}$), and activated NSCs (aNSCs: $GLAST^{+}/CD9^{high}/EGFR^{high}$) (left panel). Quantification of qNSCs, pNSCs and aNSCs populations by flow cytometry in dissociated V-SVZ cells from both genotypes (right panel). **G)** Percentage of $Ki67^{+}$, short-$BrdU^{+}$ and $ASCL1^{+}$/short-$BrdU^{+}$ cells relative to total $DAPI^{+}$ cells in the V-SVZ. **H)** Representative immunohistochemistry images showing $Ki67^{+}$ cells (green) in the V-SVZ of *Tet2-Gfap*$^{control}$ and *Tet2-Gfap*$^{cre}$mice. **I)** Quantification of $Ki67^{+}$ cells among LRC-$BrdU^{+}$ cells in the V-SVZ. **J)** Immunohistochemistry for LRC-BrdU (red) and Ki67 (green) in the V-SVZ of *Tet2-Gfap*$^{control}$ and *Tet2-Gfap*$^{cre}$ mice. **K)** Quantification of LRC-$BrdU^{+}$ cells per area in the corpus callosum (CC) of both genotypes. **L)** Representative images of LRC-$BrdU^{+}$ cells (red) in the granular cell layer (GCL) of the olfactory bulb (OB) from both genotypes (upper panel). Quantification of newborn neurons in the GL (lower panel). DAPI was used to counterstain DNA. *Gapdh* served as the normalization control for qPCR analyses. V-SVZ: ventricular-subventricular zone; LV: lateral ventricle; DIV: days *in vitro.* In bar plots, data are presented as mean ± s.e.m. In box-and-whiskers plots, the mean is indicated by (+) and whiskers denote minimun and maximum values. Statistical significance was assessed using unpaired two-tailed t-tests and Mann–Whitney tests. P-values and sample sizes (individual data points are shown) are indicated. *: $P<0.05$; ***: $P<0.001$; n.s.: not significant. Scale bars in D: 20 μm (high-magnification images, 5 μm); in H, J and L: 10 μm.

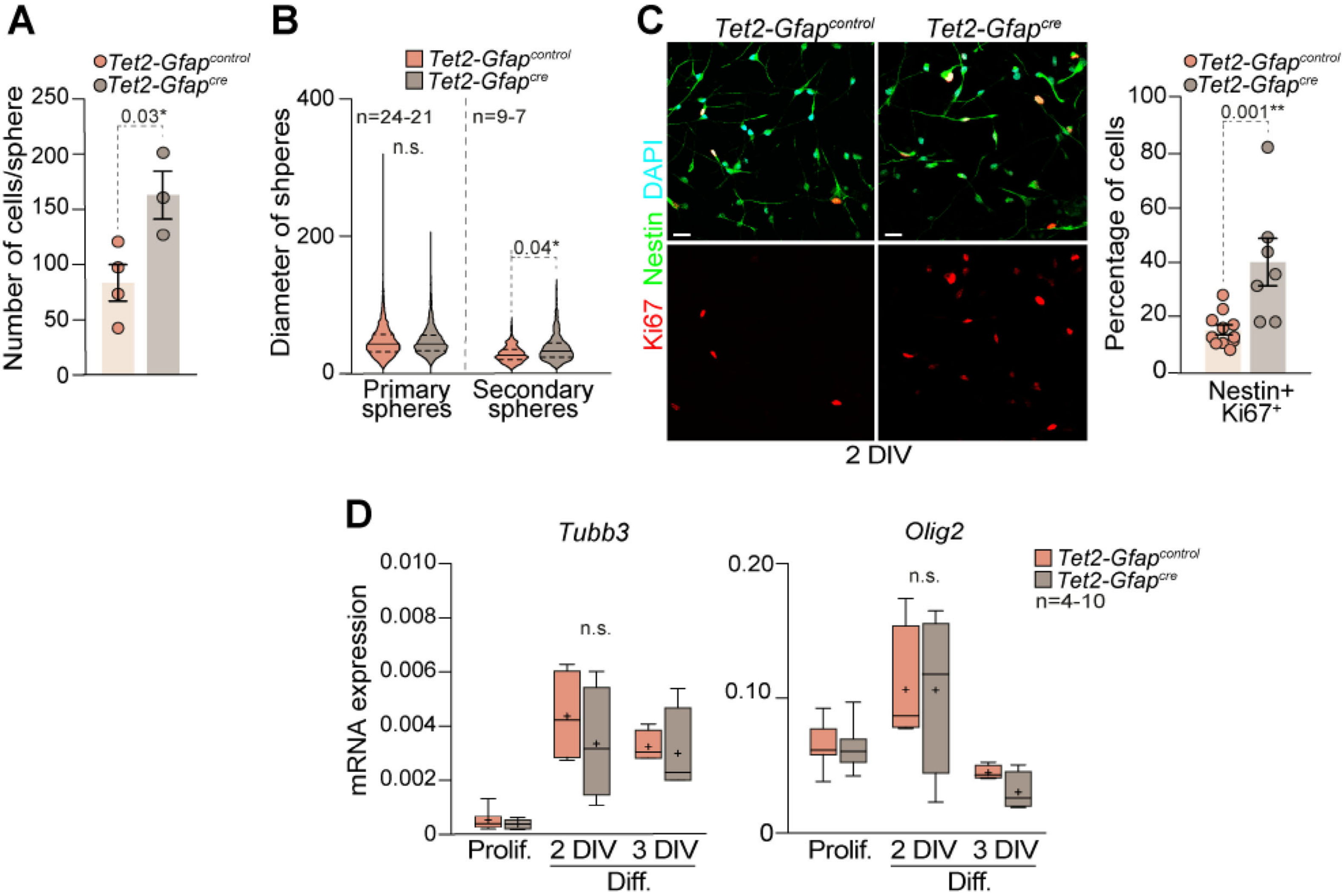


**Supplemental Figure 3. Differentiated *Tet2-Gfap^cre^* NSCs exhibit increased proliferation. Related to Figure 3. A)** Quantification of total DAPI$^+$ cells per secondary neurosphere formed from V-SVZ-derived NSCs from *Tet2-Gfap$^{control}$* and T*et2-Gfap$^{cre}$* mice. **B)** Diameters (µm) of primary and secondary spheres from NSCs cultures of both genotypes. **C)** Immunocytochemistry images of Nestin (green), and Ki67 (red) in NSCs after 2 DIV under differentiation conditions (left panel). Quantification of Nestin$^+$ and Ki67$^+$ cells relative to total DAPI$^+$ cells (right panel). **D)** qPCR analysis of differentiation markers *Tubb3* and *Olig2* in NSCs from both genotypes under proliferative (Prolif.) conditions and after 2 or 3 days of differentiation (Diff.). DAPI was used to counterstain DNA. *Gapdh* was used to normalize qPCR data. DIV: days *in vitro.* In box and whiskers plots the mean is indicated by (+) and whiskers represent the minimum and maximum values. In bar plots, data are presented as mean ± s.e.m. In violin plots, the median (dark line) and first and third quartiles (dashed lines) are shown. Statistical significance was evaluated using unpaired two-tailed t-test and Mann-Whitney test. P-values and sample sizes (n; individual data points) are indicated. *: P<0.05; **: P<0.01; n.s.: not significant. Scale bars: C, 10 µm.

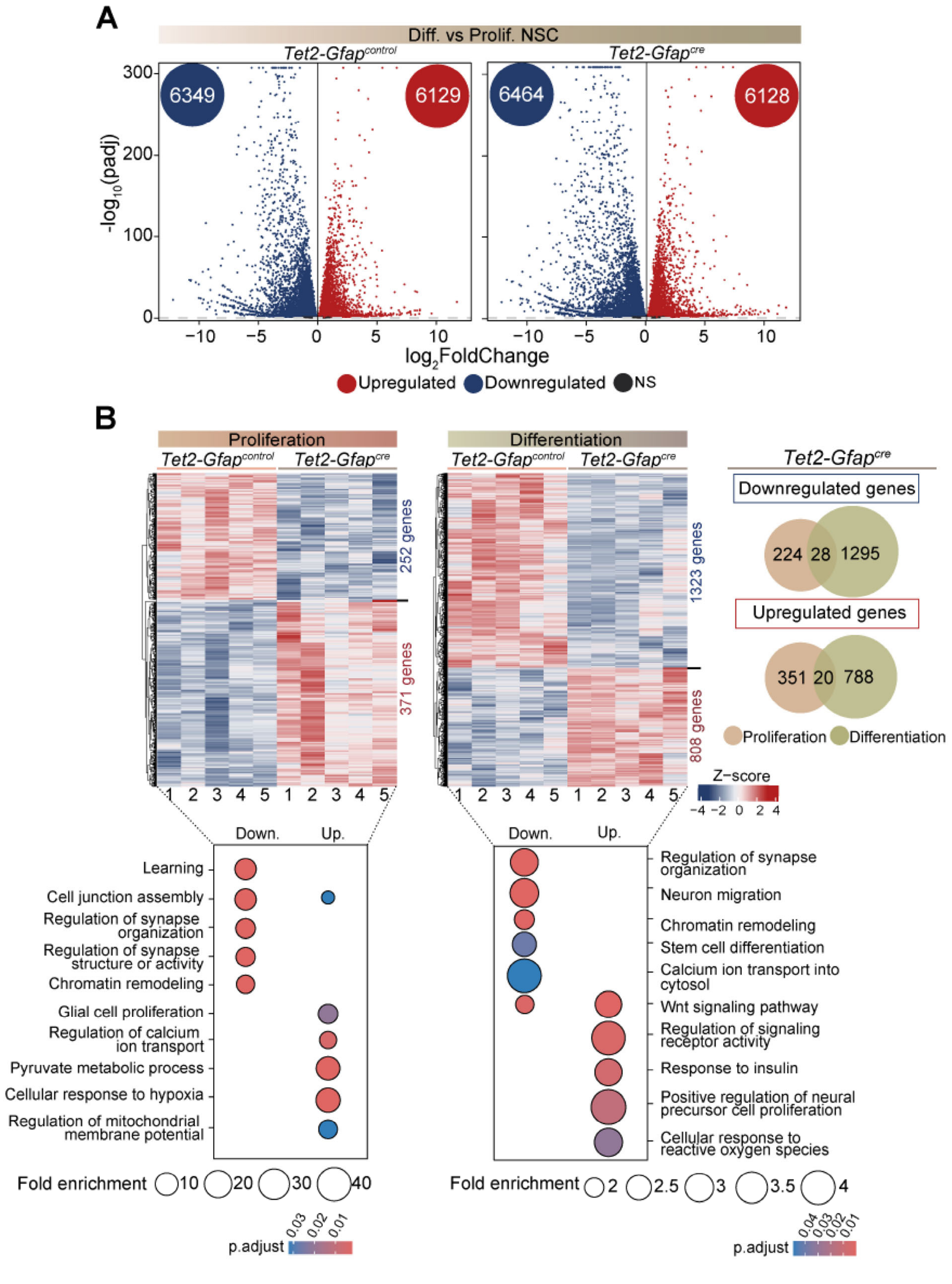


**Supplemental Figure 4. TET2 orchestrates the transcriptomic regulation of adult NSC differentiation. Related to Figure 4. A)** Volcano plots illustrating RNA-seq differential expression analysis results (log2fold change) in *Tet2-Gfap*$^{control}$ and *Tet2-Gfap*$^{cre}$ NSC cultures across the differentiation process. Upregulated genes are shown in red and downregulated genes in blue. **B)** Heatmaps showing Z-score-scaled expression of all differentially expressed genes (DEGs) comparing *Tet2-Gfap*$^{control}$ and *Tet2-Gfap*$^{cre}$ NSCs under proliferative and differentiation conditions. The number of DEGs is indicated. Venn diagrams summarizing the number of DEGs shared or uniquely regulated between proliferating and 3 DIV differentiated *Tet2-Gfap*$^{cre}$ NSCs are shown (top panel). Gene ontology (GO) enrichment analysis of biological processes associated with genes upregulated or downregulated in *Tet2-Gfap*$^{control}$ and *Tet2-Gfap*$^{cre}$ NSCs under proliferative and differentiation conditions (bottom panel).

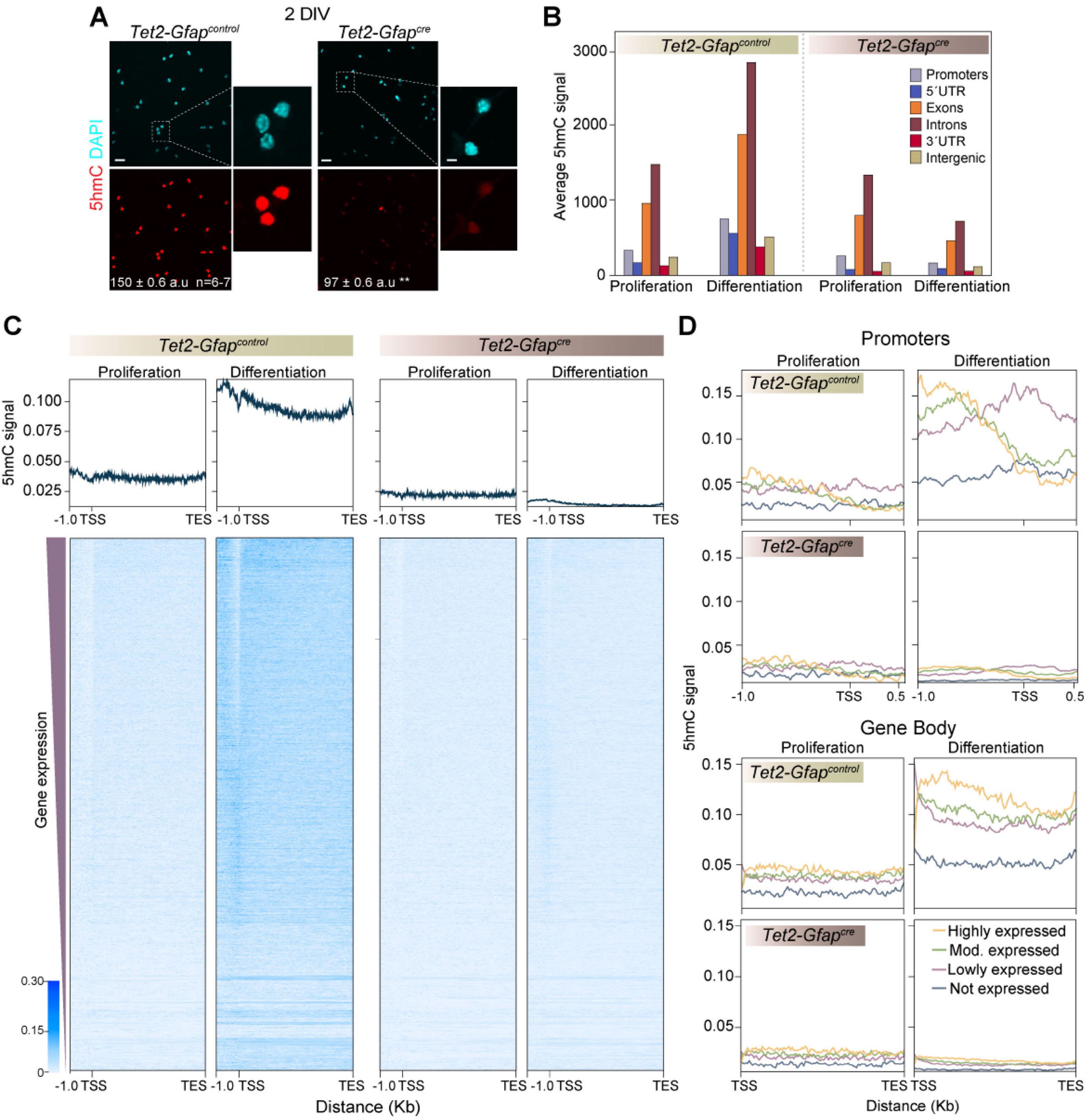


**Supplemental Figure 5. 5hmC deposition during adult NSCs differentiation is altered in the absence of *Tet2*. Related to Figure 5. A)** Immunocytochemistry images of 5hmC (red) in *Tet2-Gfap*$^{control}$ and *Tet2-Gfap*$^{cre}$ NSCs after 2 days of differentiation. Quantification of fluorescence intensity (a.u: arbitrary units) of global 5hmC levels in 2 DIV NSCs is shown. **B)** Genomic distribution of 5hmC across annotated genomic features in *Tet2-Gfap*$^{control}$ and *Tet2-Gfap*$^{cre}$ NSCs under proliferative conditions and after 3 days of *in vitro* differentiation. **C)** Heatmaps showing 5hmC enrichment from −1 kb upstream of the transcripion start site (TSS) to the transcripion end site (TES) across all annotated mouse genes, ordered by decreasing expression in 3 DIV *Tet2-Gfap*$^{control}$ NSCs. **D)** Metagene plots showing 5hmC enrichment at promoters and gene bodies, stratified by gene expression levels in NSCs of both genotypes under proliferative conditions and after 3 days of *in vitro* differentiation. DAPI was used to counterstain DNA. DIV: days *in vitro*. Significance was evaluated using unpaired two-tailed t-tests. P-values and sample sizes (n) are indicated. **: P<0.01. Scale bars in A: 10 µm (inset, 3 µm).

### Supplemental Tables

**Supplemental Table 1. List of primary antibodies used.**

| Antibody | Source | Host | Dilution | Cat # | Application |
|---|---|---|---|---|---|
| 5hmC | Active Motif | Rabbit | 1/1000 | 39769 | ICC |
| βIII-tubulin | Sigma | Rabbit | 1/300 | Ab7817 | ICC |
| ASCL1 | BD Biosciences | Mouse | 1/100 | 556604 | IHC |
| BrdU | Abcam | Rat | 1/600 | Ab6326 | IHC |
| CD31-BUV421 | BD Biosciences | Rat | 1/100 | 563356 | FC |
| CD45-BUV421 | BD Biosciences | Rat | 1/200 | 563890 | FC |
| CD9-Vio770 | Miltenyi | Rat | 1/20 | 130-102-384 | FC |
| CMS antiserum | Rao Lab | Rabbit | 1/500 | Homemade | CMS-IP |
| DCX | Abcam | Rabbit | 1/500 | Ab18723 | IHC |
| GFAP | Millipore | Chicken | 1/600 | AB5541 | ICC/IHC |
| GLAST-PE | Miltenyi | Mouse | 1/20 | 130-095-821 | FC |
| Ki67 | Invitrogen | Rat | 1/1000 | 2355034 | IHC |
| Ki67 | Abcam | Rabbit | 1/100 | ab15580 | ICC/IHC |
| Nestin | Hybridoma Bank | Mouse | 1/3 | rat-401 | ICC |
| NeuN | Millipore | Mouse | 1/50 | MAB377 | IHC |
| O4 | Cells supernatant | Mouse | 1/2 | Homemade | ICC |
| O4-405 | R&D | Rat | 1/50 | FAB1326V | FC |
| S100β | Dako | Rabbit | 1/600 | Z0311 | ICC |
| S100β | Abcam | Mouse | 1/1000 | S2532 | IHC |
| SOX2 | R&D Systems | Goat | 1:200 | AF2018 | ICC/IHC |
| Ter119-BUV421 | BD Biosciences | Rat | 1/200 | 563998 | FC |
| TET2 | ProteinTech | Rabbit | 1/100 | 21207-1-AP | ICC/IHC |

ICC, Immunocytochemistry IHC, Immunohistochemistry
FC, Flow cytometry CMS-IP, Cytosine-5-methylenesulfonate immunoprecipitation

**Supplemental Table 2. List of secondary antibodies used for immunocytochemistry.**

| Antibody | Source | Dilution | Cat # | Application |
|---|---|---|---|---|
| Alexa Fluor® 488 Donkey Anti-Chicken | Jackson ImmunoResearch | 1/600 | 703-545-155 | IHC |
| Alexa Fluor® 488 Donkey Anti-Mouse | Molecular Probes | 1/600 | A-21202 | ICC/IHC |
| Alexa Fluor® 488 Donkey Anti-Rabbit | Jackson ImmunoResearch | 1:1000 | 711-547-003 | IHC |
| Alexa Fluor® 647 Donkey Anti-Chicken | Jackson ImmunoResearch | 1/600 | 703-605-155 | ICC |
| Alexa Fluor® 647 Donkey Anti-Goat | Jackson ImmunoResearch | 1/600 | 705-605-003 | ICC/IHC |
| Alexa Fluor® 647 Donkey Anti-Rat | Jackson ImmunoResearch | 1/600 | 712-605-150 | IHC |
| Biotinylated Horse Anti-Mouse | Vector | 1/1000 | BA2000 | ICC |
| Cy3-Donkey Anti-Mouse | Jackson ImmunoResearch | 1/600 | 715-165-151 | IHC |
| Cy3-Donkey Anti-Rabbit | Jackson ImmunoResearch | 1/600 | 711-165-152 | ICC/IHC |
| Cy3-Donkey Anti-Rat | Jackson ImmunoResearch | 1/600 | 712-165-153 | IHC |
| Cy3-Streptavidin | Jackson ImmunoResearch | 1/2000 | 016-160-084 | ICC |

ICC, Immunocytochemistry IHC, Immunohistochemistry

**Supplemental Table 3. List of TaqMan probes used.** From Applied Biosystems.

| Gene | Taqman code |
|---|---|
| *Gapdh* | Mm99999915_g1 |
| *Nes* | Mm00450205_m1 |
| *Olig2* | Mm01210556_m1 |
| *S100b* | Mm00485897_m1 |
| *Tet1* | Mm01169087_m1 |
| *Tet2* | Mm00524395_m1 |
| *Tet3* | Mm00805756_m1 |
| *Tubb3* | Mm00727586_s1 |

**Supplemental Table 4. List of Syber Green primers used.**

| *Gene* | Forward (FW) | Reverse (RW) | Application |
|---|---|---|---|
| *Cre-recombinase* | GCGGTCTGGCAGTAAAAACTATC | GTGAAACAGCATTGCTGTCACTT | Expression |
| *Fgf2* | CCCACACGTCAAACTACAA | CGTCCATCTTCCTTCATAGC | Expression |
| *Gapdh* | TGGAGAAACCTGCCAAGTATG | AGTGGGAGTTGCTGTTGAAG | Expression |
| *Micu2* | GACTTGCAGAATGGCTACT | CCAAACTAATGCTCTCTCCTAC | Expression |
| *Tet2 isoform 1* | TTCTCAGAATGAAACTAGAAACTGT | TCTGGCAAACTTACATCCATTA | Expression |
| *Tet2 isoform 2* | GCAATCACCACCCAGTAGAAA | AGTACATGCTCCAAGAACAAC | Expression |
| *Tet2 genotyping* | AAGAATTGCTACAGGCCTGC | TTCTTTAGCCCTTGCTGAGC | Genotyping |
| *Tet2 exon 3* | GGTGAACAAAGTCAGAATGG | ACCCTTCTCTCCATACCTTT | Genotyping |

**Supplemental Table 5. List of software and tools used in this study.** Package name and version, and the repository for the different tools are indicated.

| *Package/Tool* | Package Version | Repository | Reference |
|---|---|---|---|
| Salmon | 1.10.1 | https://github.com/COMBINE-lab/salmon | Patro et al., 2017 |
| tximeta | 1.20.3 | https://CRAN.R-project.org/package=dplyr | Love et al., 2020 |
| DESeq2 | 1.42.1 | https://bioconductor.org/packages/DESeq2 | Love et al., 2014 |
| clusterProfiler | 4.10.1 | https://bioconductor.org/packages/clusterProfiler | T. Wu et al., 2021 |
| ggplot2 | 3.5.1 | https://cloud.r-project.org/web/packages/ggplot2/index.html | Wickham et al., 2019 |
| dplyr | 1.1.4 | https://cloud.r-project.org/web/packages/dplyr/index.html | Wickham et al., 2019 |
| tidyr | 1.3.1 | https://CRAN.R-project.org/package=tidyr | Wickham et al., 2019 |
| tibble | 3.2.1 | https://CRAN.R-project.org/package=tibble | Wickham et al., 2019 |
| ComplexHeatmap | 2.18.0 | https://bioconductor.org/packages/ComplexHeatmap | Ramírez et al., 2019 |
| Deeptols | 3.5.4 | https://github.com/deeptools/deepTools | Ramírez et al., 2019 |
| edgeR | 4.0.16 | https://bioconductor.org/packages/edgeR | Robinson et al., 2009 |
| ComBat-Ref | | https://github.com/xiaoyu12/Combat-ref | X. Zhang, 2025 |
